# Forecasting the Effects of Global Change on a Bee Biodiversity Hotspot

**DOI:** 10.1101/2024.07.10.602956

**Authors:** Mark A. Buckner, Steven T. Hoge, Bryan N. Danforth

## Abstract

The Mojave and Sonoran Deserts, recognized as a global hotspot for bee biodiversity, are experiencing habitat degradation from urbanization, utility-scale solar energy (USSE) development, and climate change. In this study, we evaluated the current and future distribution of bee diversity in the region, assessed how protected areas safeguard bee species richness, and predicted how global change may affect bees across the region. Using Joint Species Distribution Models (JSDMs) of 148 bee species, we project changes in species distributions, occurrence area, and richness across the region under four global change scenarios between 1971 and 2050. We evaluated the threat posed by USSE development and predicted how climate change will affect the suitability of protected areas for conservation. Our findings indicate that changes in temperature and precipitation do not uniformly affect bee richness across the region. Protected areas in the Sonoran and Mojave Deserts are projected to experience mean losses of up to 5.8 species, whereas protected areas at higher elevations and transition zones may gain up to 7.8 species. Outside protected areas, bee diversity is threatened by urbanization and USSE development. Areas prioritized for future USSE development have an average species richness of 4.2 species higher than the study area average, and lower priority areas have 8.2 more species. USSE zones are expected to experience declines of 2.7 to 8.0 species by 2050 due to climate change alone. Despite the importance of solitary bees for pollination, their diversity is often overlooked in land management decisions. Our results show the utility of JSDMs for extending the usability of existing data-limited bee species records, easing the inclusion of these species in conservation and land management decision-making. The multiple threats from global change drivers underscore the importance of including ecologically vital, though often data-limited, species in land-use decisions.

## Introduction

We are experiencing the onset of the sixth mass extinction (Ceballos et al., 2015). The potential for biodiversity collapse threatens ecosystem health and human welfare, which is entwined with many of the same political, economic, and social practices driving the environmental crisis itself (Sage, 2020). Humanity is responsible for the modification of 75% of the ice-free land surface (Balvanera et al., 2019; IPCC, 2022), the ballooning of atmospheric CO_2_ concentrations to levels not seen in the past two million years, and an increase in global terrestrial surface temperatures of 1.9°C over preindustrial averages (IPCC, 2023a). These global changes drive species redistributions and habitat degradation, contributing to species declines, ecosystem simplification, and a reduced capacity to provide ecosystem services (Sage, 2020).

Taking account of the looming crisis, many nations have adopted a new generation of biodiversity conservation initiatives built on the “30 by 30” framework, which seeks to protect 30% of the earth’s land and ocean area by 2030 (Convention on Biological Diversity, 2022; Dinerstein et al., 2019; European Commission & Directorate-General for Environment, 2021; Exec. Order No. 14008, 2021). The success of these initiatives depends on identifying and conserving areas of importance for biodiversity, ecosystem function, and ecosystem services. At present, over 1.2 million km^2^ are protected in the United States (US) alone (UNEP-WCMC, 2023), but these areas often have minimal utility for biodiversity conservation (Jenkins et al., 2015), instead prioritizing scenery in areas of otherwise limited economic value (Venter et al., 2018). While it is clear that we must emphasize biodiversity protection when establishing new protected areas, most assessments are taxonomically biased toward terrestrial vertebrates (Donaldson et al., 2017; Llorente-Culebras et al., 2023). Such a limited view of biodiversity is partly a result of deficient data for a substantial subset of species and ecosystems — a subset with a high proportion of threatened species (Borgelt et al., 2022).

Despite hosting relatively high biodiversity, deserts have historically received little conservation funding or research attention (Durant et al., 2012); now these same ecosystems have been designated as priority areas for utility-scale solar energy (USSE) development (Gasparatos et al., 2017). In the US, the Energy Act of 2020 increased the development pressure placed on the warm deserts of the southwestern US by establishing a minimum goal of 25 gigawatts of permitted renewable energy development by 2025 on federal lands (National Goal for Renewable Energy Production on Federal Land, 2020). While a rapid renewable energy transition is vital for addressing climate change, it may come at the cost of extensive land transformation (Capellán-Pérez et al., 2017). How this vast conversion of desert habitat for USSE will impact the ecosystem is unclear, particularly for insects (Grodsky et al., 2021; Jeal et al., 2019), which provide vital ecosystem services but often have limited demographic, distribution, and habitat preference data. These constraints mean insects may receive little consideration in the establishment of new protected areas or land management decisions.

Bees, in particular, are of public and conservation interest for the pollination services they provide and recently documented declines (Biesmeijer et al., 2006; Burkle et al., 2013; Powney et al., 2019; Turley et al., 2022). Home to upwards of a quarter of the approximately 4,000 bee species in the US, the Desert Southwest contains remarkable bee species densities (Carril et al., 2018; Minckley & Radke, 2021) making this region one of the most bee species-rich in the world (Michener, 1979; Orr et al., 2021). Many of the species documented from the southwest are host plant specialists, thought to be adapted to the variable and unpredictable desert environment (Minckley et al., 2000), and may be facing the greatest risk of decline (Bartomeus et al., 2013; Biesmeijer et al., 2006; Bogusch et al., 2020; Wood et al., 2016). While drivers of bee decline are numerous and spatially heterogeneous, many studies have focused primarily on pollinators in agroecosystems and the risks of pesticide exposure, which may be of lower importance in this region (Douglas et al., 2020). Instead, southwestern bee diversity is facing direct habitat loss from USSE, urbanization, and mineral extraction and indirect threats from introduced and invasive species (see Portman et al., 2019), and shifting disturbance regimes (Moloney et al., 2019).

The significance of climate change in bee declines remains debated (Dicks et al., 2021). As the southwest experiences more extreme temperatures (Garfin et al., 2018), increases in the duration and intensity of droughts (Williams et al., 2020), and changes in seasonal precipitation patterns (Abatzoglou & Kolden, 2011), bees may need to modify their behavior, distribution, or development to respond to heat and water stress (Johnson et al., 2023). Shifting climate may also exacerbate habitat loss by altering ecosystem composition, degrading habitat quality, and reducing floral resource availability. For example, current warming and drought conditions have already been implicated in the reduction of desert vegetation (Hantson et al., 2021) and similar patterns are expected to continue, with forecasts predicting reductions in desert perennial and forb cover and changes to shrubland composition (Munson et al., 2012).

Here, we leverage publicly available bee collection records to map the spatial distribution of bee species diversity in the deserts of the American Southwest. We predict how bee species richness changes out to 2050, to explore how bee diversity may be affected by climate-driven species redistribution and species declines. In addition, we focus on evaluating the suitability of existing protected areas and USSE development priority areas to understand the status of bee conservation and the potential threat posed by large-scale land transformation for renewable energy development and urbanization. In doing so, we argue for the consideration of understudied and data-deficient taxa in conservation and land-use decision-making through rapid and cost-effective modeling approaches.

## Methods

To explore how bee biodiversity might change under medium-term (2041-2060) global change scenarios, we utilized projected land-use change scenarios and climatology data consisting of an ensemble of eight general circulation models from the Coupled Model Intercomparison Project 6 (CMIP6) representing four shared socioeconomic pathways (SSPs; SSP1-2.6, SSP2-4.5, SSP3-7.0, and SSP5-8.5). SSPs represent greenhouse gas emissions, air pollution, and land-use trajectories from varied global socioeconomic scenarios (Riahi et al., 2017). Each of these scenarios varies in the extent of projected change from net negative emissions by 2100 and global temperature change of less than 2°C (SSP1-2.6) to a further increase in emissions through 2100 ( SSP5-8.5; IPCC, 2023b). We performed all analyses in R version 4.2.2 (R Core Team, 2022).

### Study Area

This study focused on the Mojave and Sonoran Basin and Range Omernik Level III ecoregions (Omernik, 1987). In addition, we expanded the study area to incorporate the extent of the Desert Renewable Energy Conservation Plan (DRECP), which designates priority areas for USSE development in the California deserts (BLM, 2016). To maximize the number of species that meet our minimum observation number (10 occurrences) for inclusion in our models and to reduce the risk of niche truncation by including a broader environmental gradient in our training data, we chose to buffer the study area boundary. We calculated the optimized buffer distance, *d_opt_*, by maximizing the log difference in species count per unit area as (Equation 1):

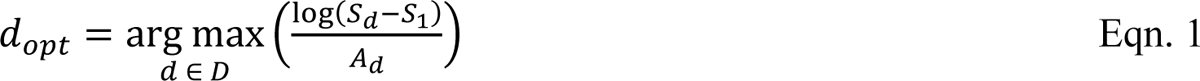

where *S_d_* is the number of species in a region with area *A_d_* associated with buffer distance *d* over a set of distances, *D*, ranging between 0 km and 48 km, and *S_1_* is the species count in the unbuffered study area. Based on data availability, we restricted the study area to the US.

### Bee Occurrence Data

We obtained occurrence records for all bee species in the Southwestern US from the Global Biodiversity Information Facility (GBIF; GBIF.org, 2023) and Symbiota Collections of Arthropods Network (SCAN; SCAN, 2022). We removed duplicate records and identified several inconsistent near duplicates between the two datasets, which had identical metadata – including unique global identifiers – but different species names. Out of caution, we excluded all records from the Bee Systematics Lab and iNaturalist entered in SCAN, prioritizing GBIF records which reported species names that were consistent with the original datasets and known from the area.

For this study, we selected a 1km grain to represent the spatial scale at which most bees interact with their environment (Kendall et al., 2022). We acknowledge that unreported or uncertain location data derived by geolocating collection records is common and our chosen grain may be smaller than the uncertainties of some records. This ‘positional certainty-ecologically informative grain’ trade-off was explored in a recent analysis which suggested that although uncertainty can reduce model performance, a coarser grain, beyond the scale at which the organism interacts with its environment, may be more detrimental (Gábor et al., 2022). To reduce uncertainty from georeferencing issues, we excluded records with a reported uncertainty greater than 5km and checked for localities corresponding to capitals, administrative area centroids, or biodiversity institutions with COORDINATECLEANER (ver. 2.0-20; Zizka et al., 2019). We compiled the final dataset by excluding any records that fell outside the buffered study area, were collected before 1971 or after 2020, had coordinates within water bodies, or represented species with less than 10 unique records.

To assess if there are any differences between how life history groups respond to climate change and urbanization, we accumulated trait data for each species regarding the degree of host plant specialization, nesting behavior (e.g., above, or below ground nest construction), life stage during winter diapause, and life history (social, solitary, or parasitic). We classified bees as host plant specialists when they were only known to collect pollen from plants within a single family or, if they were brood parasites, they parasitize a species classified as a host plant specialist (Table S1).

### Pseudo-Absence Points

The available bee occurrence records in the region are the result of opportunistic surveys without absence data and are biased toward human population centers, roadways, and certain protected areas. To facilitate modeling, we used a two-stage approach for pseudo-absence point selection. For clarity, we will distinguish between “presence-background” and “pseudo-absence” methods, though these terms are often used interchangeably (Sillero & Barbosa, 2021). Here, we characterize presence-background methods as those that use many random or biased random points to represent the full range of environmental conditions available to the species being modeled (Phillips et al., 2009). We contrast this with pseudo-absence methods which seek to generate artificial absence data (Sillero & Barbosa, 2021).

First, we calculated a normalized kernel smoothed intensity function using all available occurrence records with a 30 km bandwidth to generate a bias layer (SPATSTAT ver. 3.0-6; Baddeley et al., 2015). The resulting normalized values were treated as weights to generate 10,000 background points across the study area with similar spatial bias as the presence data (Inman et al., 2021; Kujala et al., 2015; Valavi et al., 2021). We used this initial set of background points to train balanced random forest models (BRF) for each species independently. BRF addresses class imbalance in presence-background datasets to improve predictive performance by subsampling the majority class (the background points) to equal the minority class (the presence points) at the level of each tree (Valavi et al., 2021). We trained BFR probability trees with the default parameters to estimate the occurrence probabilities of each bee species (*ranger* ver. 0.14.1; Valavi et al., 2021; Wright & Ziegler, 2017). To further refine the models, we applied recursive feature elimination, where environmental covariates were excluded based on the mean variable importance across all species. We retained the simplified models if removing a given covariate increased the area under the receiver operating characteristic (ROC) curve (AUC) for at least half of the species-level models. During each step, we estimated model performance with 5-fold spatial block cross-validation implemented with BLOCKCV (ver. 3.0-0, Valavi et al., 2019).

In the second stage, we sampled occurrence localities represented in our dataset independently for each species to use as pseudo-absence points. We weighed each record with the complement of the mean predicted species-wise occurrence probabilities from the BRF models. We chose to limit the number of background points to equal the number of presence points for each species or a maximum of 200 for honey bees (*Apis mellifera)* – a widespread species that may not be regularly collected or reported during opportunistic surveys even if it is observed.

Our method of pseudo-absence point selection allowed us to overcome several limitations of current Joint Species Distribution Model (JSDM) implementations. Notably, nearly all JSDMs require presence-absence or abundance datasets – with some exceptions in development (see Deneu et al., 2021). In presence-background methods, the number of background points should represent the full extent of available environmental conditions (Phillips et al., 2009). However, due to the considerable number of points required to do so, presence-background approaches are unfeasible, in part, because the computational cost of JSDMs scale poorly with data size (Ovaskainen & Abrego, 2020). By selecting the presence localities of other species as pseudo-absence points, we can avoid increasing the size of the dataset significantly while also ensuring that the pseudo-absence points are well informed and have similar bias as the presence points. In previous studies, related multi-step approaches outperformed random background points by directly accounting for spatial variation in sampling bias and environmental associations (Iturbide et al., 2015; Senay et al., 2013). The method we present here extends this concept from these multistep approaches to modelling contexts which scale poorly with large numbers of background points.

### Environmental Data

We considered a maximum of 18 possible environmental covariates that may affect bee distribution and habitat suitability (Table S2). Broadly, these variables represent climate (temperature and seasonal precipitation), soil sand and clay content, topography, urban land use, and forested and unforested land cover. We reprojected all environmental covariates to a custom WGS84 Albers Equal Area Conic coordinate system and resampled to 1 km resolution. Climate data were obtained from CLIMATENA (ver. 7.31), an application that downscales gridded climate data to a user-specified grain through dynamic local downscaling (Mahony et al., 2022; Wang et al., 2016). We acquired climate layers representing the average climatic conditions for each decade between 1971 and 2020 and calculated the future climate as the 10-year average of the projected annual data from CLIMATENA for 2021 through 2050 for all four SSPs. Given the medium-term time period, we assumed that the topographic and soil composition covariates will remain stable. To limit the effect of correlation on the transferability of our models, we identified all variables with an absolute Pearson’s correlation coefficient greater than 0.7 and excluded correlated variables which we expect to have lower importance for bee distributions (Feng et al., 2015). To identify any shifts in correlation structure, we checked for changes in correlation coefficients between the data period (1970 – 2020) and the future period (2021-2050) greater than ±0.1. We evaluated the extent to which extrapolation may influence our predictions under future conditions by calculating Multivariate Environmental Similarity Surfaces (MESS) for each future decade with MODEVA ver. 3.13.3 (Barbosa et al., 2013; Elith et al., 2010).

### Joint Species Distribution Model

Obtaining sufficient records to meet the suggested minimum sample sizes for single species distribution models may be impractical in community wide studies, where rare or threatened species are often a priority. To overcome this limitation, we trained JSDMs through the Hierarchical Modeling of Species Communities (HMSC) R package (HMSC ver. 3.0-14, Tikhonov et al., 2023). As opposed to stacked species distribution models, which assume each species responds independently to its environment, JSDMs assume a joint response (Norberg et al., 2019). By modeling all the species together, HMSC allows for species-environment relationships to be refined through species associations and residual variation allowing rare species to “borrow strength” from more common species (Norberg et al., 2019).

We ran four different JSDM models of varying complexity, (i) a spatial random effect only, (ii) climate covariates with no spatial random effect, (iii) all topographic and climate covariates with no spatial random effect, and (iv) all topographic and climate covariates with a spatial random effect. We used the default priors for each model and the spatial models were fit using the nearest neighbor Gaussian process (Tikhonov et al., 2020). While phylogenetic and trait data can be informative for these models, incomplete data and limited natural history observations prevented us from applying it to this analysis. For each model, we drew 500 samples from four Markov Chain Monte Carlo chains for a total of 2,000 samples. Each chain consisted of a 62,500-iteration transient period followed by sampling with a thinning interval of 250 iterations for a total of 187,500 iterations per chain. We assessed model convergence using potential scale reduction factor (Gelman & Rubin, 1992). Finally, we evaluated the explanatory power of the models with AUC and the predictive performance of these models using two-fold spatial block cross-validation (Valavi et al., 2019).

### Climate Change and Land Use

To evaluate the impacts of climate change and urbanization, we predicted the probability of occurrence for each grid cell using future climate and urbanization data for each decade and scenario. We explored the change in predicted bee richness across the study area and calculated the marginal effect of distance to the closest natural area (i.e., non-urban land cover) from the posterior sample to evaluate how the bee assemblage responds to urban sprawl. For each species, we calculated the current and future occupied area as the sum of occurrence probabilities within the study area to avoid biases from thresholding the SDM predictions (Stark & Fridley, 2022).

We then calculated range-size rarity (RSR) as a metric of grid cell importance for bee conservation in the region by identifying cells that contribute the most to the total occurrence area of range-restricted species (Guerin & Lowe, 2015). To calculate RSR without applying a threshold, we took the sum of each species occurrence probability divided by their occurrence area in each grid cell.

We evaluated how well existing protected areas overlap with predicted bee diversity by calculating the proportion of each bee’s total occurrence area within protected areas. To evaluate if protected areas differ in suitability from equivalent unprotected areas, we used the marginal effects of protected area status where all other covariates were set equal to their mean (Bakx et al., 2023). Additionally, we evaluated how the utility of individual units varies across the study area by calculating the change in average richness for each individual protected area. For this analysis, we only considered protected areas that are actively managed for conservation as reported by the US Geological Survey Protected Areas Database (USGS GAP, 2022).

Finally, we assessed how USSE development may affect bee diversity within the Southwestern US by exploring predicted species richness in areas prioritized for solar development as part of the 2012 Programmatic Environmental Impact Statement for Solar Energy Development (Solar PEIS; BLM & DOE, 2012), 2013 Restoration Design Energy Project (RDEP; BLM, 2013), and 2016 Desert Renewable Energy Conservation Plan (DRECP; BLM, 2016). We evaluated whether these current plans prioritize solar development in areas of high bee diversity and whether these areas may be of high conservation value in the future by calculating the change in average and per-unit richness in each decade and climate change scenario. Additionally, we calculated the proportion of each species occurrence area within solar development areas to identify whether certain species faced a risk substantial habitat loss from USSE development.

## Results

Our integrated and cleaned bee occurrence dataset included 5,731 records over the 5-decade data period between 1971 and 2020, where each record represents the collection of a species within a grid cell during a certain decade (see Supplemental Code). Before buffering the study area, we identified 131 species that met the minimum requirements for modeling. After buffering the study area boundary by 12 km, an additional 17 species met the modeling requirements with a median of 22 unique location-decade records each bringing the total number of modeled species to 148. Apart from honey bees (n = 1163), no species had more than 200 unique records. The final study area covers approximately 300,000 km^2,^ including portions of Arizona (AZ), California (CA), Nevada (NV), and Utah, and spans over 4,000 m in elevation (Figure 1).

**Figure 1.**
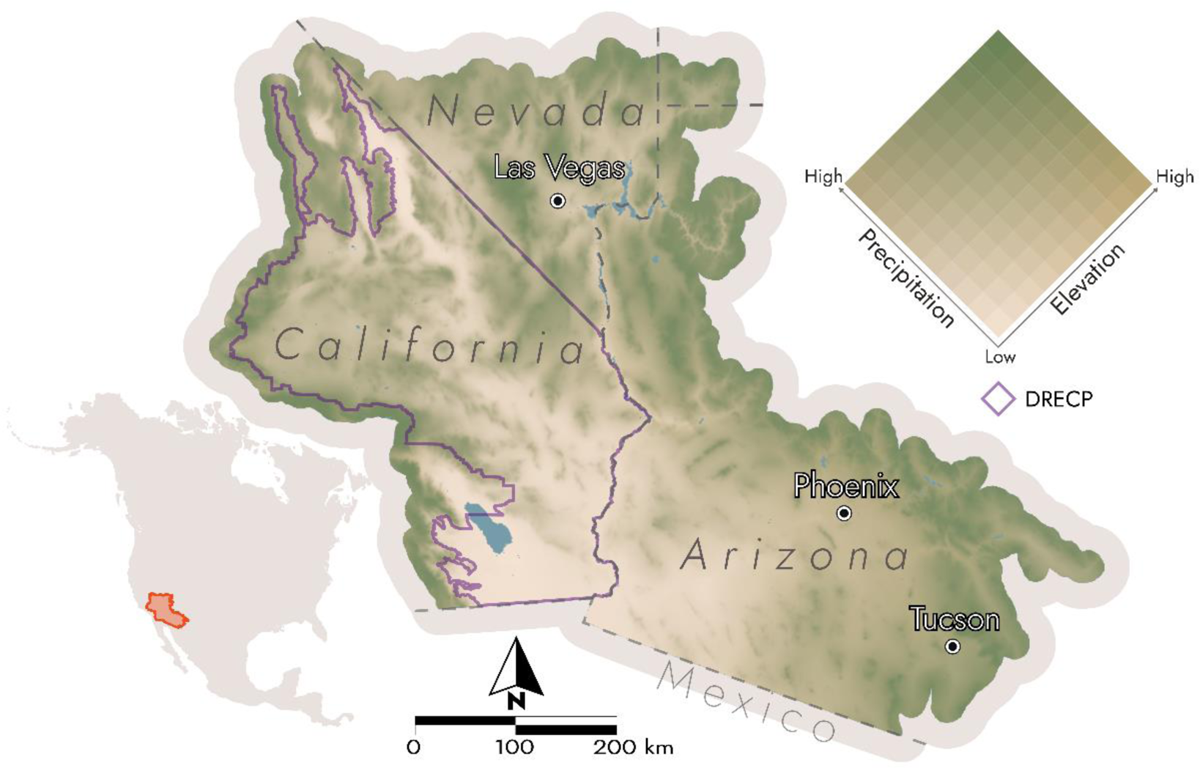
The study area covers the Mojave and Sonoran Ecoregions and the Desert Renewable Energy Conservation Plan (DRECP) area (purple) across portions of four states in the Southwestern US. The terrain color ramp corresponds to precipitation and elevation. The low, dry desert basins and valleys are colored tan, and the comparably wet mountains, plateaus, and uplands are green.

Out of our original 18 covariates, many temperature variables, including growing degree days, frost free period, and minimum temperatures, were highly correlated with extreme maximum temperature. The shift in correlation between the data period and future climate data were less than 0.1 for all pairs of covariates except for extreme maximum temperature and summer precipitation in 2050 under SSP2-4.5 which shifted by 0.12. The final covariates we chose to include in the model were extreme maximum temperature, summer and winter precipitation, annual temperature range (Wang et al., 2016), distance to natural areas within urbanized land use (Chen et al., 2022; ESA, 2017), eastness, northness, terrain ruggedness index (Amatulli et al., 2020), soil sand content (Hengl, 2018), and protected area status (USGS GAP, 2022).

### Joint Species Distribution Models

We explored fitting JSDMs with a spatial random effect, however, despite recent improvements showing promise for more efficient fitting of large spatial models (Tikhonov et al., 2020), we were not able to achieve convergence due to long computational times. Two of the four models, climate covariates only and all environmental covariates, both without spatial random effects, reached satisfactory convergence with potential scale reduction factors less than 1.1 for all parameters. Of the converged models, the simplest model performed worse than the more complex model that included both climate and topographic covariates in both explanatory and predictive performance. Overall, the more complex model performs well in explanatory tasks with an average AUC of 0.86 (range: 0.65 – 1.00). As expected, the predictive performance was lower on average, with a mean AUC of 0.67 (range: 0.05 – 1.00). We performed all the following analyses using the model with all environmental covariates and no spatial random effect.

The models suggest that climate and urbanization are the most important predictors of bee occurrence in the region. Winter precipitation accounted for a total of 24.2% of the explained variance on average, followed by distance to natural land cover with urban areas (21.5%), extreme maximum temperature (15.0%), and summer precipitation (11.5%). The remaining covariates individually accounted for no more than 7.5% of the explained variance but accounted for a substantial combined total of 27.8%. The model predicts species richness is highest in regions with moderately high extreme temperatures, peaking around 40°C and declining markedly at temperatures above this peak (Figure 2a). The effects of precipitation varied depending on the season, where higher winter precipitation resulted in lower species richness (Figure 2b). In comparison, species richness across the gradient of summer precipitation was more stable, with minor increases in richness with higher precipitation (Figure 2c). Additionally, modeled richness increased in regions with a greater difference in mean temperature between the warmest and coldest months (Figure 2d).

**Figure 2:**
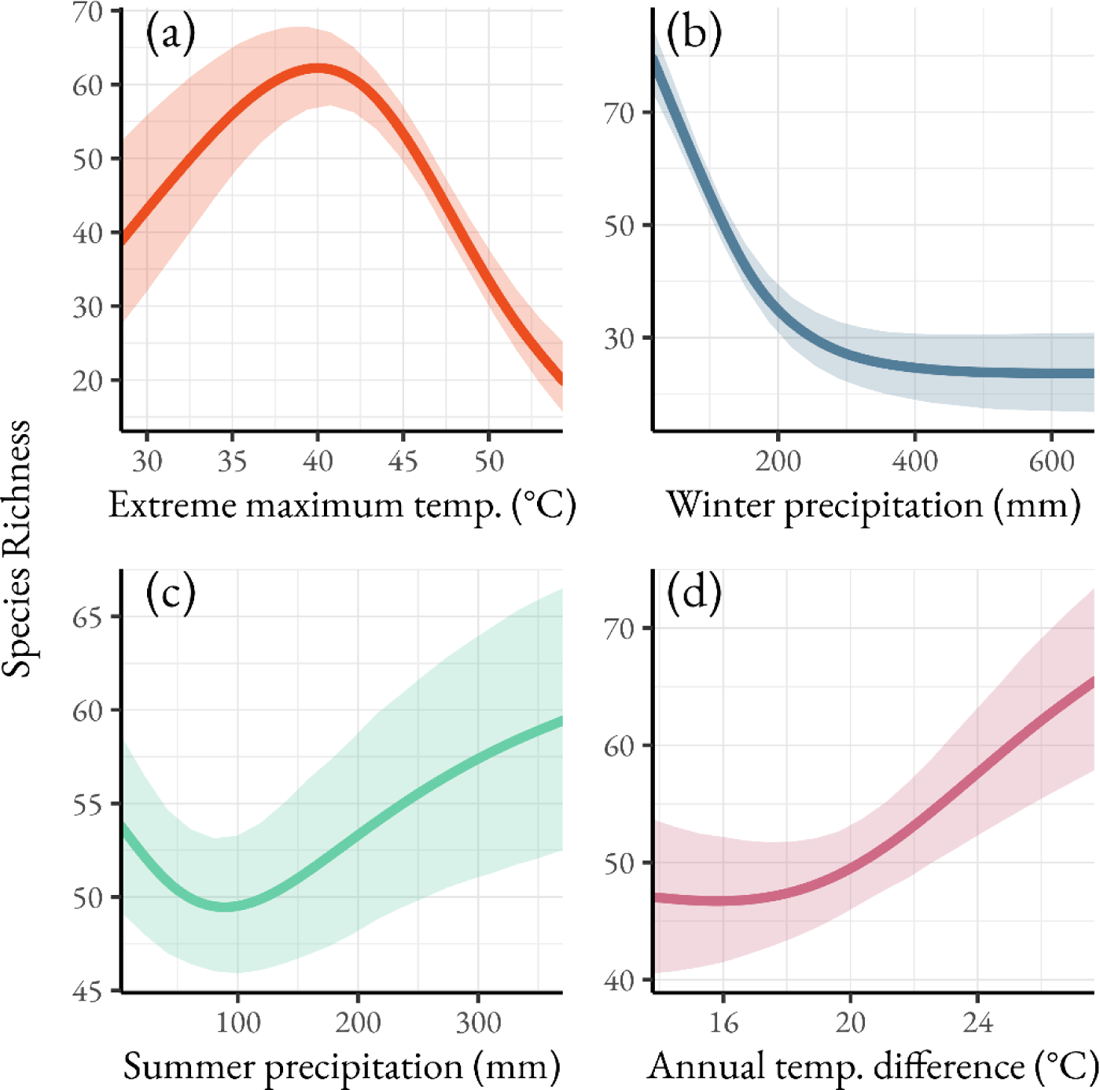
Species richness response curves to (a) extreme maximum temperature, (b) winter precipitation, (c) summer precipitation, and (d) annual temperature difference, the difference between the mean temperature during the warmest and coldest months. The shaded regions represent the 95% credible intervals.

### Climate Change and Urbanization

Species occurrence areas declined by an average of 29 km^2^ (18.5%) between 1980 and 2020. Apart from SSP 3-7.0 which continued to decline by 1.6 km^2^, our model predicts the average occurrence area will increase slightly between 2020 and 2050 by 4 km^2^ to 7.3 km^2^. The extent of habitat loss varied minimally between species grouped by life history traits. The greatest difference occurred between generalists and specialists, with generalists experiencing an average loss of only 6.5 km^2^ more than specialists. Between 23% (SSP3-7.0) and 43% (SSP2-4.5) of species are predicted to expand their ranges under climate change, with the most notable being introduced and managed honey bees (Figure S1). We did not find evidence of novel environmental conditions under future conditions, except for distance to natural areas, which required extrapolation with distances greater than 8 km (Figure S2)

During the data period, low to mid-elevations – with an exception for the greatest extremes such as Death Valley, CA – are the most important for range-restricted species (Figure 3a). RSR, the relative grid cell importance for range restricted species, declined in the lower Sonoran under future predictions and increased in the mountainous areas in the Mojave and valleys at higher latitudes and elevations, e.g., the Spring Mountains, NV, and Arizona Upland (Figure 3b). Nevertheless, the general pattern across the region remained similar to the data period, with the regions of greatest importance localized to the lower-lying areas of the Mojave and the eastern extent of the Sonoran Desert (Figure 3c, d). Beyond climate change, urbanization showed a marked effect on species richness and RSR. Within urban areas, richness declined in urban land uses as the distance to natural areas increased (Figure S3). Bees with different nesting behaviors responded differently, with the total proportion of species that nest above ground increasing by up to 1.8% in urban areas, reaching the greatest proportion of above ground nesting species at about 2.5 km from natural land cover.

**Figure 3:**
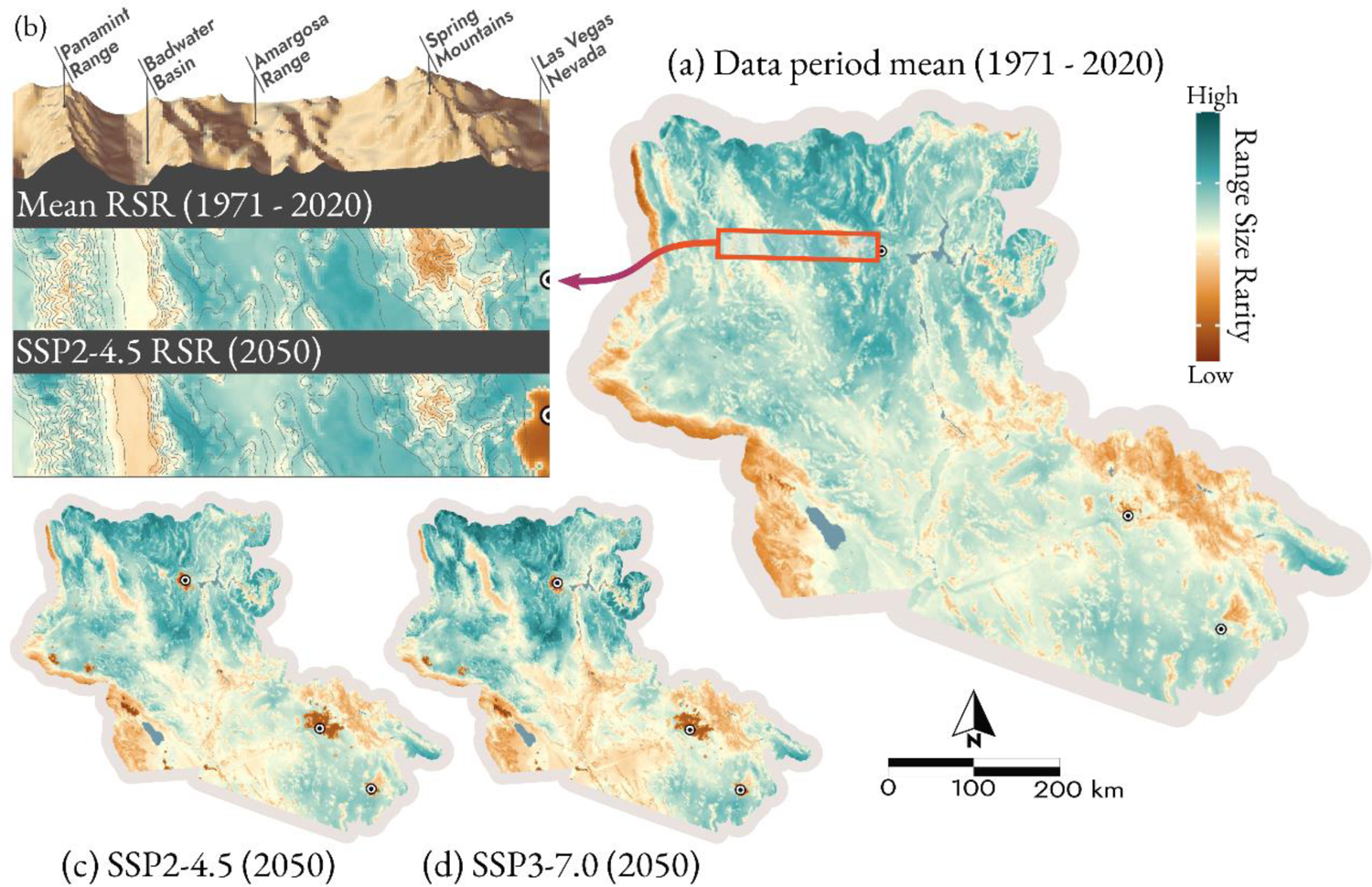
RSR is heterogeneous across the landscape (a), decreasing in low-lying valleys and urbanized areas and increasing at mid-elevations under future climate and land-use predictions (b). RSR decreases in the Sonoran between (a) the data period, calculated as the mean of five decades between 1971 and 2020, and 2050 under (c) SSP 2-4.5 and (d) SSP 3-7.0. The major metropolitan areas indicated with points are, from north to south, Las Vegas, NV; Phoenix, AZ; and Tucson, AZ.

### Protected Areas

Existing protected areas contained an average of between 21.1% and 46.1% of the predicted occurrence area for each species within the study area during the data period. The proportion of each species occurrence area contained within protected areas changed by between −1.5% for *Halictus tripartitus* to 3.7% for *Hoplitis producta* between the data period and 2050 under any SSP (Figure 4). The species with the greatest proportion of their occurrence area within protected areas have the highest occurrence probabilities at higher elevations and in the Mojave (e.g., *H. producta* and *Megandrena enceliae*), and the lowest are found primarily in the eastern Sonoran and valley floors (e.g., *Xylocopa sonorina* and *Epimelissodes duplocincta*). The marginal effects of protected areas suggest land managed under conservation mandates is no more or less suitable for any species when all other covariates are set to their mean (89% credible interval). Species richness in protected areas is predicted to decline compared to the data period average in all SSPs, with SSP 3-7.0 resulting in the greatest loss (mean = −5.8 species) and SSP 2-4.5 the lowest (mean = −2.2 species). Predicted declines vary widely across individual protected area units. By 2050, units proximal to urban areas in the Coachella Valley, CA, experience the greatest declines, losing between −27.4 species (SSP 3-7.0) and −43.8 species (SSP 5-8.5). Whereas units found within mountainous areas in and around the Mojave and the Arizona Upland increase in mean richness by up to 6.6 species in SSP 1-2.6 or 7.8 species in SSP 3-7.0 by 2050 (Figure 4). Overall, a minimum of 72% of protected land area may experience declining bee richness by 2050 with intermediate climate change (SSP 2-4.5), reaching up to 86% under SSP 3-7.0.

**Figure 4:**
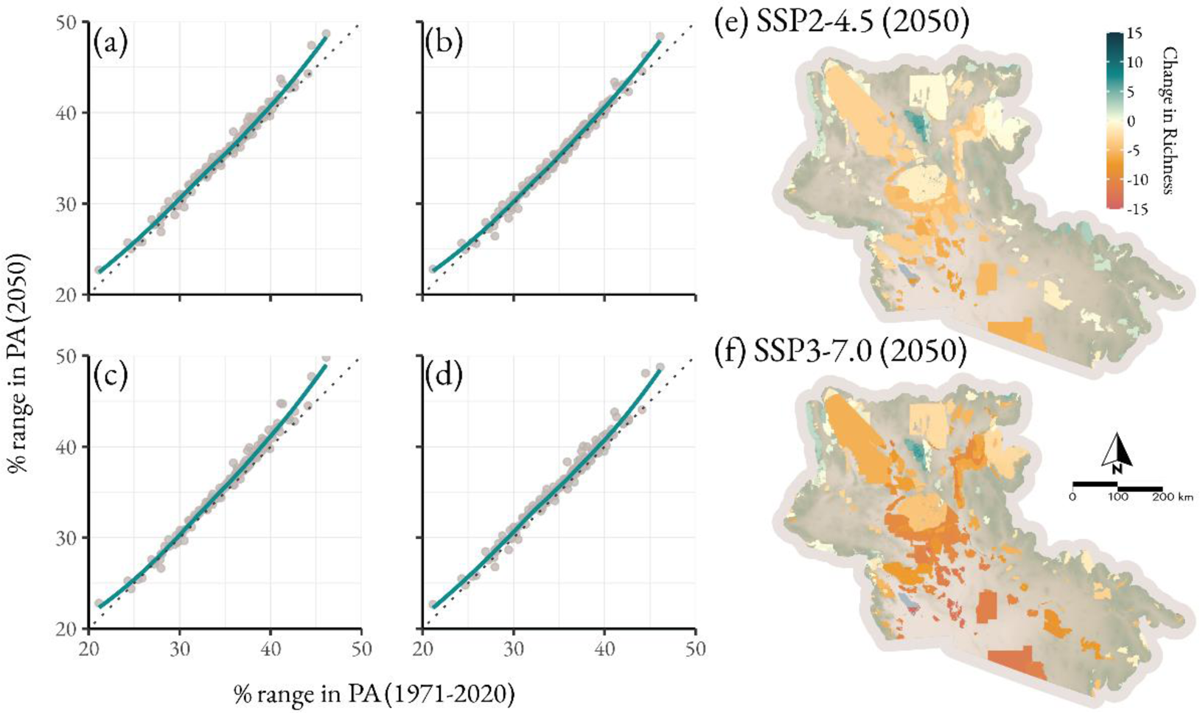
Comparison between the percent of each species occurrence area in protected areas in the data period and 2050 under (a) SSP1-2.6, (b) SSP2-4.5, (c) SSP3-7.0, and (d) SSP5-8.5. The points represent individual bee species, and the overall trend is shown with a smoothed line and dotted 1:1 line. Panels (e) and (f) show the change in per unit richness between the 1971-2020 mean and 2050 under (e) SSP2-4.5 and (f) SSP3-7.0.

### USSE Development

USSE priority and variance areas, which span over 16,000 km^2^ in the study area, contained above average species richness during the data period (priority areas: mean = 74.2 species, variance areas: mean = 78.2 species, study area: mean = 70.0 species). By 2020, the predicted richness of these areas declined to an average of 65.1 species in priority areas (variance areas: 71.8 species) after which predicted species richness did not continue to decline, remaining between 63.6 (SSP 3-7.0) and 68.4 species (SSP 5-8.5) on average. The individual responses of species vary depending on the extent to which their range overlapped with USSE development areas during the data period. For bees with the greatest proportions of their range in USSE development areas, the predicted occurrence area overlapping with USSE does not deviate substantially from the data period average. In contrast, species with less than 5.5% of their occurrence area in USSE development areas had a smaller overlap under future conditions (Figure 5). Overall, the average richness of USSE development areas is expected to decline by 2050 in all climate scenarios with the largest declines in SSP 3-7.0 (−8.0 species) with all other scenarios declining by between −2.7 (SSP 2-4.5) and −4.7 (SSP 1-2.6) species. Like protected areas, the greatest declines are expected in USSE units near urbanizing land uses with a maximum loss between −59.4 species in SSP 3-7.0 and −63.1 species in SSP 5-8.5. Notably, some units within transition zones may see an increase in richness between 4.8 species (SSP 1-2.6) and 5.3 species (SSP 2-4.5).

**Figure 5:**
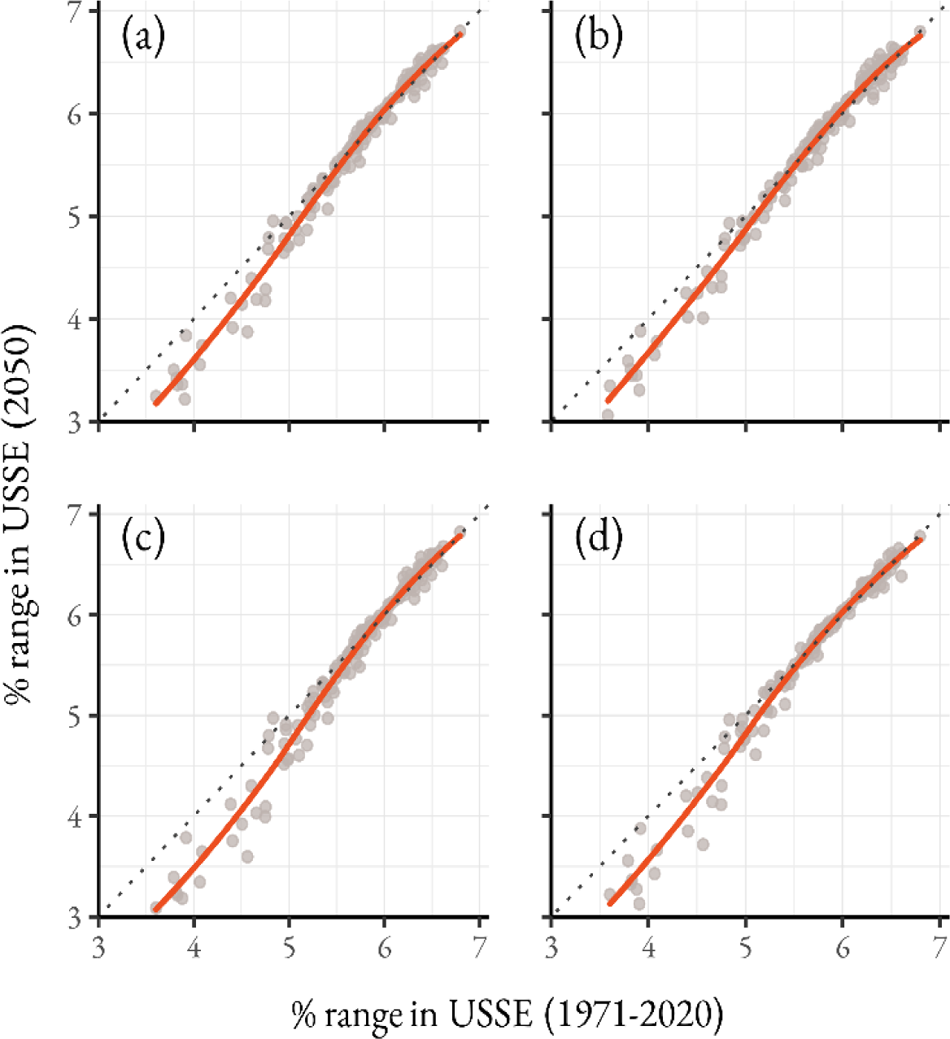
Comparison between the percent of each species range in solar energy development areas during the data period (1971-2020) and 2050 under (a) SSP1-2.6, (b) SSP2-4.5, (c) SSP3-7.0, and (d) SSP5-8.5. The points represent individual species, and the deviation of the trend from the dotted 1:1 line is shown with a smoothed line.

## Discussion

Our findings suggest that the diverse arid-adapted bee assemblage of the Mojave and Sonoran deserts is not immune to the effects of climate and land-use change. By accounting for the changing climate and extensive urbanization, our model suggests that bee richness is likely to decline across up to 90% of the Mojave and Sonoran ecoregions under midterm (2041-2060) global change, although the effects are heterogeneous across the region and climate change scenario. Current protected areas are not uniquely suited for protecting bee diversity compared to surrounding unprotected land, and on the whole existing units will continue to protect a substantial portion, at least 21%, of both current and future bee ranges, even though we expect richness to decline in many individual units.

Our model supports the assertion that warming temperatures and altered precipitation patterns will negatively affect native bee diversity in the coming decades. In the context of the few existing arid-adapted bee species distribution models, our findings, with few exceptions (e.g., honey bees), differ in predicting range contractions as opposed to expansions (Dew et al., 2019; Silva et al., 2018). Although the implications of climate change remain uncertain, explorations of thermal physiology suggest that desert bees may be vulnerable to extreme heat and desiccation, although the effects vary depending on the species and past exposure (Barrett et al., 2022; Bennett et al., 2021; Burdine & McCluney, 2019; Chappell, 1982; Hamblin et al., 2017; Johnson, Alvarez, et al., 2023). In this study, bee richness peaked at an intermediate extreme temperature of around 40°C, suggesting extreme high temperatures reduce habitat suitability for many species. Even so, bees may be able to respond through altered behavior or adaptation (Johnson, Glass, et al., 2023). Bet-hedging, in particular, could help ensure bee emergence remains synchronous with host plant bloom year to year and the most hazardous environmental conditions are avoided (Danforth, 1999). Such adaptations may prove particularly important for persistence under future climate conditions, as increasingly long duration droughts make floral resource availability more unpredictable (Minckley et al., 2013).

Beyond climate change, one of the most prominent examples of land-use change driving habitat loss in the region is the expansion of rapidly growing urban areas (Wu et al., 2011). Our models predicted that continued urban sprawl, and concurrently the infilling of habitat fragments in desert metropolises (Shrestha et al., 2012), is likely to exacerbate bee declines. Existing literature on the effects of urbanization on desert bees supports our findings. Previous work in Phoenix, AZ, found that bee richness is higher in desert habitats outside urban areas than in urban desert fragments or residential areas (Hostetler & McIntyre, 2001). In addition, our findings of a minor increase in the proportion of cavity nesting bees in urbanized areas. This pattern is consistent with past observations in the urban matrix of Tucson, AZ, that found an increase in nesting resource availability for cavity nesting bees resulted in an over representation of these species in urban fragments relative to undeveloped desert sites (Cane et al., 2006).

Within the near term, USSE is poised to contribute to rapid and extensive land-use change in the southwest. While the exact implications of solar installations on insects are not fully known, our results indicate that development is slated to occur in regions of high bee diversity. The degree to which solar installations may contribute to additional declines in bee richness will likely depend on the method of site preparation. For non-bee floral visitors, any method of site preparation, whether blading or mowing, resulted in displacement (Grodsky et al., 2021) and both methods reduce perennial plant and cacti cover, potentially limiting floral resource availability (Grodsky & Hernandez, 2020). While we were not able to directly model the effects of solar development, our results suggest that USSE priority areas contain high bee richness under both current and future climatic conditions. Even so, development is not occurring in climate refugia, and we predict developing areas to experience declining richness and lower importance for range restricted species in the future due to climate change, particularly in the Sonoran Desert. Although, we note some exceptions to this trend in units within the Arizona Upland and northern Mojave. Individual species may have different degrees of susceptibility to the co-occurring threats of development and climate change, and we emphasize that unraveling the consequences of these threats happening concurrently is a critical area of future research required to rigorously evaluate the risks posed by USSE to pollinators.

While urbanization and renewable energy development are expanding, much of the Desert Southwest is composed of protected lands that are actively managed for conservation. These protected areas currently overlap with a minimum of 21% of each bee species occurrence area, but not all units maintain high richness under mid-term predictions with over 72% of protected land area expected to experience declines. As with bee richness overall, protected areas in the Sonoran Desert and low-lying areas in the Mojave are likely to experience declining richness in the future. Units located nearest to cities showed the greatest loss of species richness in our models, but it is reasonable to equate these to large desert fragments which are still able to maintain relatively high levels of bee diversity compared to the surrounding urban matrix (Cane et al., 2006), suggesting that our model may be overpredicting species losses in this circumstance. Our results suggest an increasing importance of conserving habitats at higher elevations in the Mojave and the Arizona Uplands. The declining richness of protected areas highlights the need to reconsider establishing new conservation lands to protect climate refugia. As bees redistribute to track their climate niche, higher elevations and topographically rough landscapes could provide suitable microclimates for displaced species. Central to the concept of “conserving nature’s stage” (Beier et al., 2015), prioritizing abiotic diversity may serve to better protect southwest pollinators in the coming decades.

Our models were not able to account for the full complexity of species-ecosystem interactions under global change. Various aspects of how bees respond to climate and land-use change cannot be readily accounted for in correlative presence/pseudo-absence models, such as host plant interactions, dispersal ability, or demographics. Moreover, our models may not account for behavioral or adaptive responses to the environment which may make species less susceptible to climate change. Such uncertainty is particularly evident in situations where the future environmental conditions do not have a present-day analog requiring extrapolation into novel conditions, which we only identified with extreme distances to natural areas under future urban sprawl. We further caution that our predictions represent our best hypothesis of how future change may impact richness in the region and overall. The usefulness of AUC for presence/pseudo-absence data is limited (Golicher et al., 2012), however, it remains the most common method to measure the support of presence-only models (Konowalik & Nosol, 2021). Lastly, it is important to acknowledge that the use of pseudo-absence data itself required assumptions about where each species is least likely to occur which may introduce uncertainty in the species environment response and stochasticity in the predictions themselves. Ideally, such an analysis could be replicated many times and the results combined to generate final predictions, however, the long runtime of our models made this impractical. Additionally, null model approaches that require repeat model fitting using random draws of presence records are similarly impractical (Raes & ter Steege, 2007). Our approach using existing occurrence locations of other species means that both pseudo-absence and presence data may not represent the full environmental response for each species. While we utilized the best available data, future work in this area would benefit from additional records, ideally structured collection efforts that aim to reduce collection biases, report absence localities or non-detections, and better represent the full environmental gradient in the region.

Considering the rapid pace of development and push for the establishment of new biodiversity targets, embracing modeling methods is necessary to facilitate the consideration of invertebrate species in conservation decision-making. While the push for a structured national pollinator monitoring program is gaining ground (Woodard et al., 2020) many of the threats to pollinator diversity are allowed to move forward, as land managers and policymakers are unable to adequately consider the potential impacts due to deficient historical data. The DRECP exemplifies this need, in that the planners acknowledged the high diversity of invertebrate species within the planning area but excluded them from consideration in the planning process (BLM, 2016). The work we present here serves as an example of leveraging existing data sources to predict the distributions of rare and data-limited species. We emphasize the need to consider future climatic conditions in the establishment of protected areas and new development. Conservation actions, such as increasing native vegetation in residential areas, preserving intact habitat patches in USSE and urban developments, and protecting habitat that provide diverse microclimates that could serve as refugia are vital for conserving bee diversity and the services they provide within one of their most diverse ecosystems.

## Acknowledgements

We thank the taxonomists, ecologists, naturalists, and community scientists who collect and publicly share bee locality data. The members of the Danforth Lab and Cornell’s Pollinator Reading Group provided valuable feedback throughout this project. The Cornell Atkinson Center for Sustainability funded this work through the Sustainable Biodiversity Fund awarded to MAB.

## Data Availability Statement

The occurrence data and scripts used in this study are archived on Figshare: https://doi.org/10.6084/m9.figshare.26081812.

