## Supplemental files for "Forecasting the Effects of Global Change on a Bee Biodiversity Hotspot"

**Table S1:** Trait data for all modeled bee species

| Species | Subgenus | Sociality | Parasite | Nest | Specialist | Oil Collection | Over-wintering | Origin | Diurnality | References |
|---|---|---|---|---|---|---|---|---|---|---|
| *Agapostemon angelicus* | Agapostemon | Solitary | No | ground | No | No | Adult | Native | Diurnal | 15, 16, 19, 24 |
| *Agapostemon melliventris* | Agapostemon | Solitary | No | ground | No | No | Adult | Native | Diurnal | 15, 16, 19, 24 |
| *Ancylandrena larreae* | - | Solitary | No | ground | Yes | No | Prepupa | Native | Diurnal | 15, 16, 23, 24, 45, 46 |
| *Ancylandrena timberlakei* | - | Solitary | No | ground | No | No | Prepupa | Native | Diurnal | 15, 16, 24, 46 |
| *Andrena cerasifolii* | - | Solitary | No | ground | No | No | Adult | Native | Diurnal | 15, 16, 24 |
| *Andrena fracta* | Plastandrena | Solitary | No | ground | No | No | Adult | Native | Diurnal | 15, 16, 24, 34 |
| *Andrena palpalis* | Belandrena | Solitary | No | ground | No | No | Adult | Native | Diurnal | 15, 16, 24, 34 |
| *Andrena prunorum* | Plastandrena | Solitary | No | ground | No | No | Adult | Native | Diurnal | 15, 16, 24, 32 |
| *Andrena sphaeralceae* | - | Solitary | No | ground | Yes | No | Adult | Native | Diurnal | 15, 16, 24, 25 |
| *Anthidiellum ehrhorni* | Loyolanthidium | Solitary | No | above | No | No | Prepupa | Native | Diurnal | 15, 16, 24 |
| *Anthidiellum notatum* | Loyolanthidium | Solitary | No | above | No | No | Prepupa | Native | Diurnal | 15, 16, 24 |
| *Anthidiellum robertsoni* | Loyolanthidium | Solitary | No | above | No | No | Prepupa | Native | Diurnal | 15, 16, 24 |
| *Anthidium cockerelli* | Anthidium | Solitary | No | ground | No | No | Prepupa | Native | Diurnal | 15, 16, 24 |
| *Anthidium emarginatum* | Anthidium | Solitary | No | ground | No | No | Prepupa | Native | Diurnal | 15, 16, 24 |
| *Anthidium jocosum* | Anthidium | Solitary | No | ground | No | No | Prepupa | Native | Diurnal | 15, 16, 24 |
| *Anthidium maculosum* | Anthidium | Solitary | No | above | No | No | Prepupa | Native | Diurnal | 15, 16, 24 |
| *Anthidium palmarum* | Anthidium | Solitary | No | ground | No | No | Prepupa | Native | Diurnal | 15, 16, 24 |
| *Anthidium paroselae* | Anthidium | Solitary | No | ground | No | No | Prepupa | Native | Diurnal | 15, 16, 24 |

| Species | Subgenus | Sociality | Parasite | Nest | Specialist | Oil Collection | Over-wintering | Origin | Diurnality | References |
|---|---|---|---|---|---|---|---|---|---|---|
| *Anthophora californica* | Anthophoroides | Solitary | No | ground | No | No | Prepupa | Native | Diurnal | 15, 16, 24, 63 |
| *Anthophora centriformis* | Paramegilla | Solitary | No | ground | No | No | Prepupa | Native | Diurnal | 15, 16, 24, 63 |
| *Anthophora cockerelli* | Micranthophora | Solitary | No | ground | No | No | Prepupa | Native | Diurnal | 15, 16, 24, 63 |
| *Anthophora coptognatha* | Lophanthophora | Solitary | No | ground | No | No | Prepupa | Native | Diurnal | 15, 16, 24, 63 |
| *Anthophora curta* | Micranthophora | Solitary | No | ground | No | No | Prepupa | Native | Diurnal | 15, 16, 24, 63 |
| *Anthophora hololeuca* | Micranthophora | Solitary | No | ground | No | No | Prepupa | Native | Diurnal | 15, 16, 24, 63 |
| *Anthophora neglecta* | Lophanthophora | Solitary | No | ground | No | No | Prepupa | Native | Diurnal | 15, 16, 24, 63 |
| *Anthophora pachyodonta* | Micranthophora | Solitary | No | ground | No | No | Prepupa | Native | Diurnal | 15, 16, 24, 63 |
| *Anthophora petrophila* | Micranthophora | Solitary | No | ground | No | No | Prepupa | Native | Diurnal | 15, 16, 24, 63 |
| *Anthophora phenax* | Micranthophora | Solitary | No | ground | No | No | Prepupa | Native | Diurnal | 15, 16, 24, 63 |
| *Anthophora urbana* | Mystacanthophora | Solitary | No | ground | No | No | Prepupa | Native | Diurnal | 15, 16, 24, 63 |
| *Anthophora vannigera* | Pyganthophora | Solitary | No | ground | No | No | Prepupa | Native | Diurnal | 15, 16, 24 |
| *Apis mellifera* | Apis | Social | No | above | No | No | Adult | Non-Native | Diurnal | 15, 16, 24 |
| *Ashmeadiella bigeloviae* | Ashmeadiella | Solitary | No | above | No | No | Prepupa | Native | Diurnal | 15, 16, 24, 43 |
| *Ashmeadiella breviceps* | Arogochila | Solitary | No | above | No | No | Prepupa | Native | Diurnal | 15, 16, 24, 43 |
| *Ashmeadiella bucconis* | Ashmeadiella | Solitary | No | above | Yes | No | Prepupa | Native | Diurnal | 15, 16, 23, 24, 43 |
| *Ashmeadiella femorata* | Ashmeadiella | Solitary | No | above | No | No | Prepupa | Native | Diurnal | 15, 16, 24, 43 |
| *Ashmeadiella meliloti* | Ashmeadiella | Solitary | No | above | No | No | Prepupa | Native | Diurnal | 15, 16, 24, 43 |
| *Ashmeadiella prosopidis* | Ashmeadiella | Solitary | No | above | No | No | Prepupa | Native | Diurnal | 15, 16, 24, 43 |

| Species | Subgenus | Sociality | Parasite | Nest | Specialist | Oil Collection | Over-wintering | Origin | Diurnality | References |
|---|---|---|---|---|---|---|---|---|---|---|
| *Ashmeadiella rhodognatha* | Chilosima | Solitary | No | above | No | No | Prepupa | Native | Diurnal | 15, 16, 24, 43 |
| *Augochlorella pomoniella* | - | Social | No | ground | No | No | Adult | Native | Diurnal | 9,15, 16, 24 |
| *Bombus californicus* | Thoracobombus | Social | No | ground | No | No | Adult | Native | Diurnal | 15, 16, 24 |
| *Bombus crotchii* | Cullumanobombus | Social | No | ground | No | No | Adult | Native | Diurnal | 15, 16, 24 |
| *Bombus huntii* | Pyrobombus | Social | No | ground | No | No | Adult | Native | Diurnal | 15, 16, 24 |
| *Bombus melanopygus* | Pyrobombus | Social | No | ground | No | No | Adult | Native | Diurnal | 15, 16, 24 |
| *Bombus morrisoni* | Cullumanobombus | Social | No | ground | No | No | Adult | Native | Diurnal | 15, 16, 24 |
| *Bombus pensylvanicus* | Thoracobombus | Social | No | ground | No | No | Adult | Native | Diurnal | 15, 16, 24 |
| *Bombus sonorus* | Thoracobombus | Social | No | ground | No | No | Adult | Native | Diurnal | 15, 16, 24 |
| *Bombus vandykei* | Pyrobombus | Social | No | ground | No | No | Adult | Native | Diurnal | 15, 16, 24 |
| *Bombus vosnesenskii* | Pyrobombus | Social | No | ground | No | No | Adult | Native | Diurnal | 15, 16, 24 |
| *Brachymelecta californica* | - | Solitary | Yes | ground | No | No | Prepupa | Native | Diurnal | 15, 16, 24, 36 |
| *Calliopsis anomoptera* | Perissander | Solitary | No | ground | Yes | No | Prepupa | Native | Diurnal | 15, 16, 24, 59 |
| *Calliopsis puellae* | Nomadopsis | Solitary | No | ground | Yes | No | Prepupa | Native | Diurnal | 15, 16, 24, 59 |
| *Calliopsis rozeni* | Calliopsima | Solitary | No | ground | Yes | No | Prepupa | Native | Diurnal | 15, 16, 24, 59 |
| *Calliopsis subalpina* | Hypomacrotera | Solitary | No | ground | Yes | No | Prepupa | Native | Diurnal | 15, 16, 24, 59 |
| *Centris atripes* | Paracentris | Solitary | No | ground | No | Yes | Prepupa | Native | Diurnal | 15, 16, 20, 24 |
| *Centris cockerelli* | Paracentris | Solitary | No | ground | No | Yes | Prepupa | Native | Diurnal | 15, 16, 20, 24 |
| *Centris hoffmanseggiae* | Paracentris | Solitary | No | ground | No | No | Prepupa | Native | Diurnal | 15, 16, 20, 24 |

| Species | Subgenus | Sociality | Parasite | Nest | Specialist | Oil Collection | Over-wintering | Origin | Diurnality | References |
|---|---|---|---|---|---|---|---|---|---|---|
| *Centris pallida* | Paracentris | Solitary | No | ground | No | No | Prepupa | Native | Diurnal | 8, 15, 16, 20, 24 |
| *Centris rhodopus* | Paracentris | Solitary | No | ground | No | Yes | Prepupa | Native | Diurnal | 15, 16, 20, 24 |
| *Ceratina apacheorum* | Zadontomerus | Solitary | No | above | No | No | Adult | Native | Diurnal | 15, 16, 24 |
| *Ceratina arizonensis* | Ceratinula | Solitary | No | above | No | No | Adult | Native | Diurnal | 15, 16, 24, 41 |
| *Chelostoma californicum* | Neochelostoma | Solitary | No | above | Yes | No | Prepupa | Native | Diurnal | 15, 16, 24 |
| *Colletes cercidii* | - | Solitary | No | ground | No | No | Prepupa | Native | Diurnal | 15, 16, 24, 60, 61 |
| *Colletes clypeonitens* | - | Solitary | No | ground | Yes | No | Prepupa | Native | Diurnal | 15, 16, 24 |
| *Colletes larreae* | - | Solitary | No | ground | Yes | No | Prepupa | Native | Diurnal | 15, 16, 23, 24 |
| *Colletes louisae* | - | Solitary | No | ground | No | No | Prepupa | Native | Diurnal | 15, 16, 24 |
| *Colletes salicicola* | - | Solitary | No | ground | No | No | Prepupa | Native | Diurnal | 15, 16, 24 |
| *Colletes stepheni* | - | Solitary | No | ground | Yes | No | Prepupa | Native | Nocturnal | 15, 14, 16, 24 |
| *Conanthalictus bakeri* | - | Solitary | No | ground | Yes | No | Prepupa | Native | Diurnal | 15, 16, 24, 52 |
| *Conanthalictus caerulescens* | - | Solitary | No | ground | Yes | No | Prepupa | Native | Diurnal | 15, 16, 24, 52 |
| *Diadasia australis* | Coquillettapis | Solitary | No | ground | Yes | No | Prepupa | Native | Diurnal | 15, 16, 24 |
| *Diadasia bituberculata* | Coquillettapis | Solitary | No | ground | Yes | No | Prepupa | Native | Diurnal | 15, 16, 24 |
| *Diadasia diminuta* | Coquillettapis | Solitary | No | ground | Yes | No | Prepupa | Native | Diurnal | 15, 16, 24 |
| *Diadasia lutzi* | Coquillettapis | Solitary | No | ground | Yes | No | Prepupa | Native | Diurnal | 15, 16, 24 |
| *Diadasia martialis* | Coquillettapis | Solitary | No | ground | Yes | No | Prepupa | Native | Diurnal | 15, 16, 24 |
| *Diadasia rinconis* | Coquillettapis | Solitary | No | ground | Yes | No | Prepupa | Native | Diurnal | 15, 16, 24, 33,37 |

| Species | Subgenus | Sociality | Parasite | Nest | Specialist | Oil Collection | Over-wintering | Origin | Diurnality | References |
|---|---|---|---|---|---|---|---|---|---|---|
| *Diadasia tuberculifrons* | Coquillettapis | Solitary | No | ground | Yes | No | Prepupa | Native | Diurnal | 15, 16, 24, 33 |
| *Diadasia vallicola* | Coquillettapis | Solitary | No | ground | Yes | No | Prepupa | Native | Diurnal | 15, 16, 24, 33 |
| *Dianthidium dubium* | Dianthidium | Solitary | No | above | No | No | Prepupa | Native | Diurnal | 15, 16, 24 |
| *Dianthidium pudicum* | Dianthidium | Solitary | No | above | No | No | Prepupa | Native | Diurnal | 15, 16, 24 |
| *Dianthidium ulkei* | Dianthidium | Solitary | No | above | No | No | Prepupa | Native | Diurnal | 15, 16, 21, 24 |
| *Dieunomia nevadensis* | Epinomia | Solitary | No | ground | No | No | Prepupa | Native | Diurnal | 11, 15, 16, 24, 67 |
| *Dioxys productus* | - | Solitary | Yes | above | No | No | Prepupa | Native | Diurnal | 15, 16, 24 |
| *Dufourea mulleri* | - | Solitary | No | ground | Yes | No | Prepupa | Native | Diurnal | 15, 16, 24, 66 |
| *Epeolus mesillae* | - | Solitary | Yes | ground | Yes | No | Prepupa | Native | Diurnal | 15, 16, 24, 35 |
| *Ericrocis lata* | - | Solitary | Yes | ground | No | No | Prepupa | Native | Diurnal | 15, 16, 24, 49 |
| *Eucera mohavensis* | Synhalonia | Solitary | No | ground | No | No | Prepupa | Native | Diurnal | 15, 16, 17, 24, 62 |
| *Habropoda pallida* | - | Solitary | No | ground | Yes | No | Prepupa | Native | Diurnal | 2, 15, 16, 24 |
| *Habropoda tristissima* | - | Solitary | No | ground | No | No | Prepupa | Native | Diurnal | 2, 15, 16, 24 |
| *Halictus farinosus* | Nealictus | Social | No | ground | No | No | Adult | Native | Diurnal | 15, 16, 24, 30 |
| *Halictus ligatus* | Odontalictus | Social | No | ground | No | No | Adult | Native | Diurnal | 15, 16, 24, 30 |
| *Halictus tripartitus* | Seladonia | Social | No | ground | No | No | Adult | Native | Diurnal | 15, 16, 24, 30, 38 |
| *Hesperapis fuchsi* | Panurgomia | Solitary | No | ground | Yes | No | Prepupa | Native | Diurnal | 15, 16, 24 |
| *Hesperapis larreae* | Amblyapis | Solitary | No | ground | Yes | No | Prepupa | Native | Diurnal | 15, 16, 23, 24, 53 |
| *Hexepeolus rhodogyne* | - | Solitary | Yes | ground | Yes | No | Prepupa | Native | Diurnal | 15, 16, 24, 45 |

| Species | Subgenus | Sociality | Parasite | Nest | Specialist | Oil Collection | Over-wintering | Origin | Diurnality | References |
|---|---|---|---|---|---|---|---|---|---|---|
| *Hoplitis biscutellae* | Alcidamea | Solitary | No | above | Yes | No | Prepupa | Native | Diurnal | 15, 16, 24, 27 |
| *Hoplitis producta* | Alcidamea | Solitary | No | above | No | No | Prepupa | Native | Diurnal | 15, 16, 24, 27 |
| *Hylaeus asininus* | Paraprosopis | Solitary | No | above | No | No | Prepupa | Native | Diurnal | 15, 16, 24, 64 |
| *Hylaeus episcopalis* | Prosopis | Solitary | No | above | No | No | Prepupa | Native | Diurnal | 15, 16, 24 |
| *Hylaeus wootoni* | Paraprosopis | Solitary | No | above | No | No | Prepupa | Native | Diurnal | 15, 16, 24 |
| *Lasioglossum hyalinum* | Dialictus | Social | No | ground | No | No | Adult | Native | Diurnal | 15, 16, 24, 56 |
| *Lasioglossum microlepoides* | Dialictus | Social | No | ground | No | No | Adult | Native | Diurnal | 15, 16, 24, 56 |
| *Lasioglossum perparvum* | Dialictus | Social | No | ground | No | No | Adult | Native | Diurnal | 15, 16, 24, 56 |
| *Lasioglossum pseudotegulare* | Dialictus | Social | No | ground | No | No | Adult | Native | Diurnal | 15, 16, 24, 56 |
| *Lasioglossum sisymbrii* | Lasioglossum | Solitary | No | ground | No | No | Adult | Native | Diurnal | 15, 16, 24, 56 |
| *Lasioglossum stictaspis* | Dialictus | Social | No | ground | No | No | Adult | Native | Diurnal | 15, 16, 24, 56 |
| *Lithurgopsis apicalis* | - | Solitary | No | above | No | No | Prepupa | Native | Diurnal | 15, 16, 24, 47 |
| *Lithurgopsis echinocacti* | - | Solitary | No | above | No | No | Prepupa | Native | Diurnal | 15, 16, 24, 47 |
| *Macrotera arcuata* | Macroteropsis | Solitary | No | ground | Yes | No | Prepupa | Native | Diurnal | 13, 15, 16, 24 |
| *Macrotera mellea* | Macroterella | Solitary | No | ground | No | No | Prepupa | Native | Diurnal | 13, 15, 16, 24 |
| *Megachile chilopsidis* | Chelostomoides | Solitary | No | above | No | No | Prepupa | Native | Diurnal | 3, 15, 16, 24, 28 |
| *Megachile discorhina* | Chelostomoides | Solitary | No | above | No | No | Prepupa | Native | Diurnal | 15, 16, 24, 28 |
| *Megachile fucata* | Megachiloides | Solitary | No | above | No | No | Prepupa | Native | Diurnal | 15, 16, 24, 28 |
| *Megachile* | Litomegachile | Solitary | No | above | No | No | Prepupa | Native | Diurnal | 15, 16, 24, |

| Species | Subgenus | Sociality | Parasite | Nest | Specialist | Oil Collection | Over-wintering | Origin | Diurnality | References |
|---|---|---|---|---|---|---|---|---|---|---|
| *lippiae* | | | | | | | | | | 28 |
| *Megachile newberryae* | Sayapis | Solitary | No | above | No | No | Prepupa | Native | Diurnal | 15, 16, 24, 28 |
| *Megachile odontostoma* | Chelostomoides | Solitary | No | above | No | No | Prepupa | Native | Diurnal | 15, 16, 24 , 28 |
| *Megachile policaris* | Sayapis | Solitary | No | above | No | No | Prepupa | Native | Diurnal | 15, 16, 24, 28 |
| *Megachile sidalceae* | Pseudocentron | Solitary | No | above | No | No | Prepupa | Native | Diurnal | 15, 16, 24, 28 |
| *Megandrena enceliae* | Megandrena | Solitary | No | ground | Yes | No | Prepupa | Native | Diurnal | 15, 16, 24 |
| *Melissodes paroselae* | Melissodes | Solitary | No | ground | No | No | Prepupa | Native | Diurnal | 15, 16, 24, 40 |
| *Melissodes tristis* | Eumelissodes | Solitary | No | ground | No | No | Prepupa | Native | Diurnal | 15, 16, 24, 40 |
| *Neolarra californica* | - | Solitary | Yes | ground | Yes | No | Prepupa | Native | Diurnal | 15, 16, 23, 24 |
| *Nomia tetrazonata* | Acunomia | Solitary | No | ground | No | No | Prepupa | Native | Diurnal | 15, 16, 24, 67 |
| *Osmia aglaia* | Melanosmia | Solitary | No | above | No | No | Adult | Native | Diurnal | 4, 15, 16, 24, 55 |
| *Osmia ribifloris* | Osmia | Solitary | No | above | No | No | Adult | Native | Diurnal | 4, 15, 16, 24 |
| *Perdita albonotata* | Procockerellia | Solitary | No | ground | Yes | No | Prepupa | Native | Diurnal | 12, 15, 16, 24, 31 |
| *Perdita arenaria* | Heteroperdita | Solitary | No | ground | Yes | No | Prepupa | Native | Diurnal | 15, 16, 24, 31 |
| *Perdita callicerata* | Hexaperdita | Solitary | No | ground | Yes | No | Prepupa | Native | Diurnal | 15, 16, 24, 31 |
| *Perdita coldeniae* | Heteroperdita | Solitary | No | ground | Yes | No | Prepupa | Native | Diurnal | 15, 16, 24, 31 |
| *Perdita covilleae* | Perdita | Solitary | No | ground | Yes | No | Prepupa | Native | Diurnal | 15, 16, 24, 31 |
| *Perdita koebelei* | Perdita | Solitary | No | ground | Yes | No | Prepupa | Native | Diurnal | 15, 16, 24, 31 |
| *Perdita larreae* | Perditella | Solitary | No | ground | Yes | No | Prepupa | Native | Diurnal | 15, 16, 23, 24, 31 |

| Species | Subgenus | Sociality | Parasite | Nest | Specialist | Oil Collection | Over-wintering | Origin | Diurnality | References |
|---|---|---|---|---|---|---|---|---|---|---|
| *Perdita lateralis* | Perdita | Solitary | No | ground | Yes | No | Prepupa | Native | Diurnal | 15, 16, 24, 31 |
| *Perdita malacothricis* | Pygoperdita | Solitary | No | ground | Yes | No | Prepupa | Native | Diurnal | 15, 16, 24, 31 |
| *Perdita minima* | Perditella | Solitary | No | ground | Yes | No | Prepupa | Native | Diurnal | 15, 16, 24, 31 |
| *Perdita mohavensis* | Pygoperdita | Solitary | No | ground | Yes | No | Prepupa | Native | Diurnal | 15, 16, 24, 31 |
| *Perdita punctosignata* | Perdita | Solitary | No | ground | Yes | No | Prepupa | Native | Diurnal | 15, 16, 24, 31 |
| *Perdita punctulata* | Perdita | Solitary | No | ground | Yes | No | Prepupa | Native | Diurnal | 15, 16, 24, 31 |
| *Pseudomacrotera turgiceps* | - | Solitary | No | ground | Yes | No | Prepupa | Native | Diurnal | 15, 16, 24 |
| *Protodufourea eickworti* | - | Solitary | No | ground | Yes | No | Prepupa | Native | Diurnal | 15, 16, 24, 54 |
| *Protohalonia amoena* | - | Solitary | No | ground | Yes | No | Prepupa | Native | Nocturnal | 14, 15, 16, 24 |
| *Protosmia rubifloris* | Chelostomopsis | Solitary | No | above | No | No | Adult | Native | Diurnal | 15, 16, 22, 24 |
| *Protoxaea gloriosa* | - | Solitary | No | ground | No | No | Prepupa | Native | Diurnal | 15, 16, 24 |
| *Stelis perpulchra* | Dolichostelis | Solitary | Yes | above | No | No | Prepupa | Native | Diurnal | 15, 16, 24 |
| *Epimelissodes duplocincta* | Idiomelissodes | Solitary | No | ground | Yes | No | Prepupa | Native | Diurnal | 15, 16, 24 |
| *Epimelissodes sabinensis* | Epimelissodes | Solitary | No | ground | No | No | Prepupa | Native | Diurnal | 15, 16, 24 |
| *Townsendiella pulchra* | - | Solitary | Yes | ground | Yes | No | Prepupa | Native | Diurnal | 15, 16, 24, 53 |
| *Trachusa larreae* | Heteranthidium | Solitary | No | ground | Yes | No | Prepupa | Native | Diurnal | 15, 16, 23, 24, 50 |
| *Triepeolus verbesinae* | - | Solitary | Yes | ground | No | No | Adult | Native | Diurnal | 15, 16, 24 |
| *Xeralictus bicuspidariae* | - | Solitary | No | ground | No | No | Prepupa | Native | Diurnal | 15, 16, 24, 42 |
| *Xeralictus timberlakei* | - | Solitary | No | ground | No | No | Prepupa | Native | Diurnal | 15, 16, 24, 42 |

| Species | Subgenus | Sociality | Parasite | Nest | Specialist | Oil Collection | Over-wintering | Origin | Diurnality | References |
|---|---|---|---|---|---|---|---|---|---|---|
| *Xylocopa californica* | Xylocopoides | Solitary | No | above | No | No | Prepupa | Native | Diurnal | 1, 15, 16, 24 |
| *Xylocopa sonorina* | Neoxylocopa | Solitary | No | above | No | No | Prepupa | Native | Diurnal | 1, 15, 16, 24, 58 |
| *Xylocopa tabaniformis* | Notoxylocopa | Solitary | No | above | No | No | Prepupa | Native | Diurnal | 1, 15, 16, 24 |

**Sociality**, all species exhibiting a degree of social behavior from eusociality to cooperative breeding are marked social. Communally nesting bees are considered solitary. **Parasite** indicates species which parasitize nests of other bee species. **Nest** categorizes species into two broad groups- those that make nests below ground (ground) and those that nest above ground (above) including species nesting in cavities, stems, or which construct above ground nests. **Specialist** indicates species recorded to only collect pollen from one plant family are classified as specialists. **Oil collection** indicates if species is known to collect floral oils. **Overwintering** indicates if a species is known to overwinter as an adult or prepupa. **Origin**, whether the bee is native to the Southwest. **Diurnality**, both nocturnal and crepuscular bees were considered nocturnal. Trait data were inferred for species without sufficient natural history records based on taxonomy.

**Table S1 References:**


1. Ackerman, A. J. (1916). The carpenter-bees of the United States of the genus Xylocopa. *Journal of the New York Entomological Society*, *24*(3), 196-232.
2. Alcock, J., & Buchmann, S. (2011). The mating system of Habropoda pallida Timberlake (Anthophorinae: Apidae). Journal of insect behavior, 24, 348-362.
3. Armbrust, E. A. (2004). Resource use and nesting behavior of Megachile prosopidis and M. chilopsidis with notes on M. discorhina (Hymenoptera: Megachilidae). *Journal of the Kansas Entomological Society*, *77*(2), 89-98.
4. Bosch, J., Maeta, Y., & Rust, R. (2001). A phylogenetic analysis of nesting behavior in the genus Osmia (Hymenoptera: Megachilidae). *Annals of the Entomological Society of America*, *94*(4), 617-627.
5. Bossert, S., Wood, T. J., Patiny, S., Michez, D., Almeida, E. A., Minckley, R. L., ... & Murray, E. A. (2022). Phylogeny, biogeography and diversification of the mining bee family Andrenidae. Systematic Entomology, 47(2), 283-302.
6. Burks, B. D. (1968). The pollen-collecting bees of the Anthidiini of California (Hymenoptera: Megachilidae).
7. Cane, J. H., & Neff, J. L. (2011). Predicted fates of ground-nesting bees in soil heated by wildfire: thermal tolerances of life stages and a survey of nesting depths. Biological Conservation, 144(11), 2631-2636.
8. Chappell, M. A. (1984). Temperature regulation and energetics of the solitary bee Centris pallida during foraging and intermale mate competition. *Physiological Zoology*, *57*(2), 215-225.
9. Correia, M. (1980). Study of the biology of Heriades truncorum. 1. Biological and morphological aspects. *Apidologie*, *11*(4), 309-339.
10. Cross, E. A., & Bohart, G. E. (1960). The biology of Nomia (Epinomia) triangulifera with comparative notes on other species of Nomia. University of Kansas Science Bulletin, 41(6), 761.
11. Danforth, B. N. (1989). Nesting behavior of four species of Perdita (Hymenoptera: Andrenidae). *Journal of the Kansas Entomological Society*, 59-79.
12. Danforth, B. N., Ji, S., & Ballard, L. J. (2003). Gene flow and population structure in an oligolectic desert bee, Macrotera (Macroteropsis) portalis (Hymenoptera: Andrenidae). *Journal of the Kansas Entomological Society*, 221-235.
13. Danforth, B. N., Minckley, R. L., & Neff, J. L. (2019). The solitary bees: biology, evolution, conservation. Princeton University Press.
14. Diller, S.N., Schaeffer, J.S., Grundel, R., Pavlovic, N.B., McKenna, J.E. Jr., Esselman, P.C., 2020, Bee-Gap: Ecology, Life-History, and Distribution of Bee Species in the United States 2017: U.S. Geological Survey data release, https://doi.org/10.5066/P9QHQNNS.
15. Discoverlife, Ascher J.S., Pickering J. (2017) Discover Life bee species guide and world checklist (Hymenoptera: Apoidea: Anthophila).
16. Dorchin, A., López-Uribe, M. M., Praz, C. J., Griswold, T., & Danforth, B. N. (2018). Phylogeny, new generic-level classification, and historical biogeography of the Eucera complex (Hymenoptera: Apidae). Molecular Phylogenetics and Evolution, 119, 81-92.
17. Dyer, F. C., & Seeley, T. D. (1991). Nesting behavior and the evolution of worker tempo in four honey bee species. *Ecology*, *72*(1), 156-170.
18. Eickwort, G. C. (1981). Aspects of the nesting biology of five Nearctic species of Agapostemon (Hymenoptera: Halictidae). *Journal of the Kansas Entomological Society*, 337-351.
19. Fox, W. J. (1899). Synopsis of the United States species of the hymenopterous genus

Centris Fabr. with description of a new species from Trinidad. *Proceedings of the Academy of Natural Sciences of Philadelphia*, *51*(1), 63-70.

20. Frohlich, D. R., & Parker, F. D. (1985). Observations on the nest-building and reproductive behavior of a resin-gathering bee: Dianthidium ulkei (Hymenoptera: Megachilidae). *Annals of the Entomological Society of America*, *78*(6), 804-810.
21. Griswold, T. L. (1986). Notes on the nesting biology of Protosmia (Chelostomopsis) rubifloris (Cockerell). *The Pan-Pacific Entomologist*, *62*, 84.
22. Hurd,Paul D., Jr. and Linsley, E. Gorton. 1975. *The principal Larrea bees of the southwestern United States (Hymenoptera, Apoidea)*. Washington: Smithsonian Institution Press. In *Smithsonian Contributions to Zoology*, 193. https://doi.org/10.5479/si.00810282.193.
23. Krombein, K. V., Hurd, P. D., Smith, D. R., & Burks, B. D. (1979). *Catalog of Hymenoptera in America north of Mexico*(Vol. 1, pp. 1199-2209). Washington, DC: Smithsonian Institution Press.
24. LaBerge, W. E. (1969). A revision of the bees of the genus Andrena of the Western Hemisphere. Part II. Plastandrena, Aporandrena, Charitandrena. *Transactions of the American Entomological Society (1890-)*, *95*(1), 1-47.
25. LaBerge, W. E. (2001). Revision of the bees of the genus Tetraloniella in the New World (Hymenoptera: Apidae). *Illinois Natural History Survey Bulletin; v. 036, no. 03*.
26. Michener, C. D. (1947). A revision of the American species of Hoplitis (Hymenoptera, Megachilidae). Bulletin of the AMNH; v. 89, article 4.
27. Michener, C. D. (1953). The biology of a leafcutter bee (Megachile brevis) and its associates. *University of Kansas Science Bulletin*, *35*(3), 1659.
28. Michener, C. D. (1975). Nests of Paranthidium jugatorium in association with Melitoma taurea (Hymenoptera: Megachilidae and Anthophoridae). *Journal of the Kansas Entomological Society*, 194-200.
29. Michener, C. D., & Bennett, F. D. (1977). Geographical variation in nesting biology and social organization of Halictus ligatus. *The University of Kansas Science Bulletin*, *51*(7), 233.
30. Michener, C. D., & Ordway, E. (1963). The life history of Perdita maculigera maculipennis (Hymenoptera: Andrenidae). *Journal of the Kansas Entomological Society*, *36*(1), 34-45.
31. Miliczky, E. (2008). Observations on the nesting biology of Andrena (Plastandrena) prunorum Cockerell in Washington state (Hymenoptera: Andrenidae). *Journal of the Kansas Entomological Society*, *81*(2), 110-121.
32. Neff, J. L., & Simpson, B. B. (1992). Partial bivoltinism in a ground-nesting bee: the biology of Diadasia rinconis in Texas (Hymenoptera, Anthophoridae). *Journal of the Kansas Entomological Society*, 377-392.
33. Neff, J. L., & Simpson, B. B. (1997). Nesting and foraging behavior of Andrena (Callandrena) rudbeckiae Robertson (Hymenoptera: Apoidea: Andrenidae) in Texas. *Journal of the Kansas Entomological Society*, 100-113.
34. Onuferko, T. M. (2019). A review of the cleptoparasitic bee genus Epeolus Latreille, 1802 (Hymenoptera: Apidae) in the Caribbean, Central America and Mexico. *European Journal of Taxonomy*, (563).
35. Onuferko, T. M., Packer, L., & Genaro, J. A. (2021). Brachymelecta Linsley, 1939, previously the rarest North American bee genus, was described from an aberrant specimen and is the senior synonym for Xeromelecta Linsley, 1939. *European Journal of Taxonomy*, *754*, 1-51.
36. Ordway, E. (1987). The life history of Diadasia rinconis Cockerell (hymenoptera:

Anthophoridae). *Journal of the Kansas Entomological Society*, 15-24.

37. Packer, L., Gravel, A. I. D., & Lebuhn, G. (2007). Phenology and social organization of Halictus (Seladonia) tripartitus (Hymenoptera: Halictidae). *Journal of Hymenoptera Research*, *16*, 281-292.
38. Parker, F. D., & Bohart, G. E. (1979). Dolichostelis, a new genus of parasitic bees (Hymenoptera: Megachilidae). *Journal of the Kansas Entomological Society*, 138-153.
39. Parker, F. D., Tepedino, V. J., & Bohart, G. E. (1981). Notes on the biology of a common sunflower bee, Melissodes (Eumelissodes) agilis Cresson. *Journal of the New York Entomological Society*, 43-52.
40. Rehan, S. M., & Richards, M. H. (2010). Nesting biology and subsociality in Ceratina calcarata (Hymenoptera: Apidae). *The Canadian Entomologist*, *142*(1), 65-74.
41. Richards, M. H., & Packer, L. (2010). Social behaviours in solitary bees: interactions among individuals in Xeralictus bicuspidariae Snelling (Hymenoptera: Halictidae: Rophitinae). *J Hym Res*, *19*, 66-76.
42. Rozen Jr, J. G. (1987). Nesting biology of the bee Ashmeadiella holtii and its cleptoparasite, a new species of Stelis (Apoidea: Megachilidae). *American Museum Novitates*, (2900).
43. Rozen Jr, J. G. (1989). Life History Studies of the "Prinmitive" Panurgine Bees (Hymenoptera: Andrenidae: Panurginae).
44. Rozen Jr, J. G. (1992). Biology of the bee Ancylandrena larreae (Andrenidae, Andreninae) and its cleptoparasite Hexepeolus rhodogyne (Anthophoridae, Nomadinae): with a review of egg deposition in the Nomadinae (Hymenoptera, Apoidea). American Museum novitates; no. 3038.
45. Rozen Jr, J. G. (1994). Biologies of the bee genera Ancylandrena (Andrenidae, Andreninae) and Hexepeolus (Apidae, Nomadinae): and phylogenetic relationships of Ancylandrena based on its mature larva (Hymenoptera, Apoidea). American Museum novitates; no. 3108.
46. Rozen Jr, J. G. (2013). Larval development and nesting biology of the adventive wood-nesting bee Lithurgus (L.) chrysurus Fonscolombe (Hymenoptera: Megachilidae: Lithurgini). *American Museum Novitates*, *2013*(3774), 20-40.
47. Rozen Jr, J. G. (2016). The Bee Svastra sabinensis: Nesting Biology, Mature Oocyte, Postdefecating Larva, and Association with Triepeolus penicilliferus (Apidae: Apinae: Eucerini and Nomadinae: Epeolini). *American Museum Novitates*, *2016*(3850), 1-12.
48. Rozen Jr, J. G., & Buchmann, S. L. (1990). Nesting biology and immature stages of the bees Centris caesalpiniae, C. pallida, and the cleptoparasite Ericrocis lata (Hymenoptera, Apoidea, Anthophoridae). American Museum novitates;; no. 2985.
49. Rozen Jr, J. G., & Hall, H. G. (2012). Nesting biology and immatures of the oligolectic bee Trachusa larreae (Apoidea: Megachilidae: Anthidiini). *American Museum Novitates*, *2012*(3765), 1-24.
50. Rozen JR, J. G., & Macneill, C. D. (1957). Biological observations on Exomalopsis (Anthophorula) chionura Cockerell, including a comparison of the biology of Exomalopsis with that of other anthophorid groups (Hymenoptera: Apoidea). *Annals of the Entomological Society of America*, *50*(5), 522-529.
51. Rozen Jr, J. G., & McGinley, R. J. (1976). Biology of the bee genus Conanthalictus (Halictidae, Dufoureinae). American Museum novitates; no. 2602.
52. Rozen Jr, J. G., & McGinley, R. J. (1991). Biology and larvae of the cleptoparasitic bee Townsendiella pulchra and nesting biology of its host Hesperapis larreae (Hymenoptera, Apoidea). American Museum novitates; no. 3005.
53. Rozen Jr, J. G., Roig-Alsina, A., & Alexander, B. A. (1997). The cleptoparasitic bee

genus Rhopalolemma: with reference to other Nomadinae (Apidae), and biology of its host Protodufourea (Halictidae, Rophitinae). American Museum novitates; no. 3194.
54. Rust, R. W. (1990). Spatial and temporal heterogeneity of pollen foraging in Osmia lignaria propinqua (Hymenoptera: Megachilidae). *Environmental Entomology*, *19*(2), 332-338.
55. Sakagami, S. F., Hoshikawa, K., & Fukuda, H. (1984). Overwintering ecology of two social halictine bees, Lasioglossum duplex and L. problematicum. *Researches on population ecology*, *26*(2), 363-378.
56. Schwarz, H. F. (1926). *North American Dianthidium, Anthidiellum, and Paranthidium*. American Museum of Natural History.
57. Sheffield, C., Heron, J., & Musetti, L. (2020). Xylocopa sonorina Smith, 1874 from Vancouver, British Columbia, Canada (Hymenoptera: Apidae, Xylocopinae) with comments on its taxonomy. *Biodiversity Data Journal*, *8*.
58. Shinn, A. F. (1967). A revision of the bee genus Calliopsis and the biology and ecology of C. andreniformis (Hymenoptera, Andrenidae).
59. Sommeijer, M. J., Neve, J., & Jacobusse, C. (2012). The typical development cycle of the solitary bee Colletes halophilus. *entomologische berichten*, *72*(1-2), 52-58.
60. Stephen, W. P. (1952). *A revision of the genus Colletes in America North of Mexico (Hymenoptera, Colletidae)* (Doctoral dissertation, University of Kansas).
61. Tepedino, V. J., & Griswold, T. L. (1995). *The bees of the Columbia Basin*. Interior Columbia Basin Ecosystem Management Project.
62. Thorp, R. W. (1969). Ecology and behavior of Anthophora edwardsii (Hymenoptera: Anthophoridae). *American Midland Naturalist*, 321-337.
63. Torchio, P. F. (1984). The nesting biology of Hylaeus bisinuatus Forster and development of its immature forms (Hymenoptera: Colletidae). *Journal of the Kansas Entomological Society*, 276-297.
64. Torchio, P. F., & Trostle, G. E. (1986). Biological notes on Anthophora urbana urbana and its parasite, Xeromelecta californica (Hymenoptera: Anthophoridae), including descriptions of late embryogenesis and hatching. *Annals of the Entomological Society of America*, *79*(3), 434-447.
65. Torchio, P. F., Rozen Jr, J. G., Bohart, G. E., & Favreau, M. S. (1967). Biology of Dufourea and of its cleptoparasite, Neopasites (Hymenoptera: Apoidea). *Journal of the New York Entomological Society*, 132-146.
66. Wcislo, W. T. (1993). Communal nesting in a North American pearly-banded bee, Nomia tetrazonata, with notes on nesting behavior of Dieunomia heteropoda (Hymenoptera: Halictidae: Nomiinae). *Annals of the entomological Society of America*, *86*(6), 813-821.

**Table S2:** Environmental covariates considered for inclusion in the JSDMs.

| Abbr. | Variable | *1971-2000* | *2001-2020* | *2021 - 2050* | Selected | Justification |
|---|---|---|---|---|---|---|
| **Climate** | | | | | | |
| EXT | Extreme Maximum Temperature[1,2] | Decade (1971-2000) | Decade (2001-2020) | Decade mean of annual data (2021 - 2100) | ✓ | Extreme heat may contribute to thermal and desiccation stress[3], and changes to floral community[4] |
| EMT | Extreme Minimum Temperature[1,2] | Decade (1971-2000) | Decade (2001-2020) | Decade mean of annual data (2021 - 2100) | | Warmer winter temperature can increase metabolic demands during overwintering[5] |
| DD1040 | Degree-days above 10°C and below 40°C[1,2] | Decade (1971-2000) | Decade (2001-2020) | Decade mean of annual data (2021 - 2100) | | Approximates flight period, growing season duration, and may relate to winter mortality[6] |
| bFFP | Beginning of Frost Free Period[1,2] | Decade (1971-2000) | Decade (2001-2020) | Decade mean of annual data (2021 - 2100) | | Possible early boundary for flight period in spring emerging species |
| eFFP | End of Frost Free Period[1,2] | Decade (1971-2000) | Decade (2001-2020) | Decade mean of annual data (2021 - 2100) | | Possible late boundary for flight period in late summer/fall emerging species |
| PPT_wt | Winter Precipitation[1,2] | Decade (1971-2000) | Decade (2001-2020) | Decade mean of annual data (2021 - 2100) | ✓ | Important determinate of bloom[7,8] |
| PPT_sm | Summer Precipitation[1,2] | Decade (1971-2000) | Decade (2001-2020) | Decade mean of annual data (2021 - 2100) | ✓ | Important determinate of bloom[8] |
| AI | Aridity Index[1,2] $AI = MAP/Eref$ | Decade (1971-2000) Derived from MAP and Eref | Decade (2001-2020) Derived from MAP and Eref | Decade mean of annual data (2021 - 2100) Derived from MAP and Eref | | Correlated with species richness in global analyses[9] |
| MAP | Mean Annual Precipitation[1,2] | Decade (1971-2000) | Decade (2001-2020) | Decade mean of annual data (2021 - 2100) | | Used in calculation of AI |

| Abbr. | Variable | *1971-2000* | *2001-2020* | *2021 - 2050* | Selected | Justification |
|---|---|---|---|---|---|---|
| MAT | Mean Annual Temperature[1,2] | Decade (1971-2000) | Decade (2001-2020) | Decade mean of annual data (2021 - 2100) | ✓ | Warming temperatures may cause thermal stress and higher metabolic demands during overwintering[3,5] |
| Eref | Hargreaves Reference Evapotranspiration[1,2] | Decade (1971-2000) | Decade (2001-2020) | Decade mean of annual data (2021 - 2100) | | Used in calculation of AI |
| **Soils** | | | | | | |
| SSC | Soil sand content at 0cm[10] | Assumed fixed | Assumed fixed | Assumed fixed | ✓ | Plausible contributor to nest site selection in ground nesting species. May also be a lose proxy for plant community.[11] |
| SCC | Soil clay content at 0cm[12] | Assumed fixed | Assumed fixed | Assumed fixed | | Plausible contributor to nest site selection in ground nesting species. May also be a lose proxy for plant community.[11] |
| **Topography** | | | | | | |
| Eness | Eastness[13] | Assumed fixed | Assumed fixed | Assumed fixed | ✓ | Alters nest site suitability and solar exposure[11] |
| Nness | Northness[13] | Assumed fixed | Assumed fixed | Assumed fixed | ✓ | Alters nest site suitability and solar exposure[11] |
| TRI | Terrain Ruggedness Index[13] | Assumed fixed | Assumed fixed | Assumed fixed | ✓ | Proxy for microclimate availability and habitat heterogeneity[14] |
| CTI | Compound Topographic Index[13] | Assumed fixed | Assumed fixed | Assumed fixed | | Proxy for microclimate availability and habitat heterogeneity[14] |
| **Land Use** | | | | | | |
| LULC | Land Use, Land Cover[15,16] (Simplified to forested and | Note: For 1971 - 1990 used data for 1992 | **2010:** Used ESA[6] data **2020:** Projected data | **All periods and SSPs:** End points (e.g., for 2020 - 2030 use | | Different land use types support varied floral and bee communities |

| Abbr. | Variable | *1971-2000* | *2001-2020* | *2021 - 2050* | Selected | Justification |
|---|---|---|---|---|---|---|
| | unforested cover) | | | 2025) data | | |
| ULF | Urban Land Fraction[17–19] | Historical Data: 1975 to 20148 Switched to projected data source for 2000 | Ten-year intervals of urban grid cell fraction for 2010 and 2020 | Ten-year intervals of urban grid cell fraction for 2030 to 2050 | | Urbanization linked to changes in bee diversity and community compositions[20,21] |
| PA | Protected Areas[22] | Assumed fixed | Assumed fixed | Assumed fixed | ✓ | Included to facilitate analysis |
| DNA | Distance to Natural Area | Derived from LULC | Derived from LULC | Derived from LULC | ✓ | Urban fringe has more available habitat due to lower density development[23] |

**Table S2 References:**


1. Wang, T., Hamann, A., Spittlehouse, D. & Carroll, C. Locally downscaled and spatially customizable climate data for historical and future periods for North America. *PLoS ONE* **11**, (2016).
2. Mahony, C. R., Wang, T., Hamann, A. & Cannon, A. J. A global climate model ensemble for downscaled monthly climate normals over North America. *International Journal of Climatology* **42**, 5871–5891 (2022).
3. Johnson, M. G., Glass, J. R., Dillon, M. E. & Harrison, J. F. How will climatic warming affect insect pollinators? in *Advances in Insect Physiology* vol. 64 1–115 (Elsevier, 2023).
4. Moloney, K. A. *et al.* Increased fire risk in Mojave and Sonoran shrublands due to exotic species and extreme rainfall events. *Ecosphere* **10**, e02592 (2019).
5. Williams, C. M., Henry, H. A. L. & Sinclair, B. J. Cold truths: how winter drives responses of terrestrial organisms to climate change. *Biological Reviews* **90**, 214–235 (2015).
6. Sgolastra, F. *et al.* The long summer: Pre-wintering temperatures affect metabolic expenditure and winter survival in a solitary bee. *Journal of Insect Physiology* **57**, 1651–1659 (2011).
7. Beatley, J. C. Phenological Events and Their Environmental Triggers in Mojave Desert Ecosystems. *Ecology* **55**, 856–863 (1974).
8. Bowers, J. E. Has Climatic Warming Altered Spring Flowering Date of Sonoran Desert Shrubs? *The Southwestern Naturalist* **52**, 347–355 (2007).
9. Orr, M. C. *et al.* Global Patterns and Drivers of Bee Distribution. *Current Biology* **31**, 451-458.e4 (2021).
10. Hengl, T. Sand content in % (kg / kg) at 6 standard depths (0, 10, 30, 60, 100 and 200 cm) at 250 m resolution. Zenodo https://doi.org/10.5281/zenodo.2525662 (2018).
11. Antoine, C. M. & Forrest, J. R. K. Nesting habitat of ground-nesting bees: a review. *Ecological Entomology* **46**, 143–159 (2021).
12. Hengl, T. Clay content in % (kg / kg) at 6 standard depths (0, 10, 30, 60, 100 and 200 cm) at 250 m resolution. Zenodo https://doi.org/10.5281/ZENODO.1476854 (2018).
13. Amatulli, G., McInerney, D., Sethi, T., Strobl, P. & Domisch, S. Geomorpho90m, empirical evaluation and accuracy assessment of global high-resolution geomorphometric layers. *Sci Data* **7**, 162 (2020).
14. Title, P. O. & Bemmels, J. B. ENVIREM: an expanded set of bioclimatic and topographic variables increases flexibility and improves performance of ecological niche modeling. *Ecography* **41**, 291–307 (2018).
15. ESA. Land Cover CCI Product User Guide Version 2. Tech. Rep. (2017).
16. Chen, G., Li, X. & Liu, X. Global land projection based on plant functional types with a 1-km resolution under socio-climatic scenarios. *Sci Data* **9**, 125 (2022).
17. Corbane, C., Florczyk, A., Pesaresi, M., Politis, P. & Syrris, V. GHS-BUILT R2018A - GHS built-up grid, derived from Landsat, multitemporal (1975-1990-2000-2014) - OBSOLETE RELEASE. European Commission, Joint Research Centre (JRC) https://doi.org/10.2905/jrc-ghsl-10007 (2018).
18. Gao, J. & Pesaresi, M. Downscaling SSP-consistent global spatial urban land projections from 1/8-degree to 1-km resolution 2000–2100. *Sci Data* **8**, 281 (2021).
19. Gao, J. & Pesaresi, M. Global 1-km Downscaled Urban Land Extent Projection and Base Year Grids by SSP Scenarios, 2000-2100. NASA Socioeconomic Data and Applications Center (SEDAC) (2021).
20. Hostetler, N. E. & McIntyre, M. E. Effects of urban land use on pollinator (Hymenoptera: Apoidea) communities in a desert metropolis. *Basic and Applied Ecology* **2**, 209–218 (2001).
21. Cane, J. H., Minckley, R. L., Kervin, L. J., Roulston, T. H. & Williams, N. M. Complex Responses Within A Desert Bee Guild (Hymenoptera: Apiformes) To Urban Habitat Fragmentation. *Ecological Applications* **16**, 632–644 (2006).

22. USGS GAP. Protected Areas Database of the United States (PAD-US). U.S. Geological Survey data release https://doi.org/10.5066/P9Q9LQ4B (2022).
23. Shrestha, M. K., York, A. M., Boone, C. G. & Zhang, S. Land fragmentation due to rapid urbanization in the Phoenix Metropolitan Area: Analyzing the spatiotemporal patterns and drivers. *Applied Geography* **32**, 522–531 (2012).

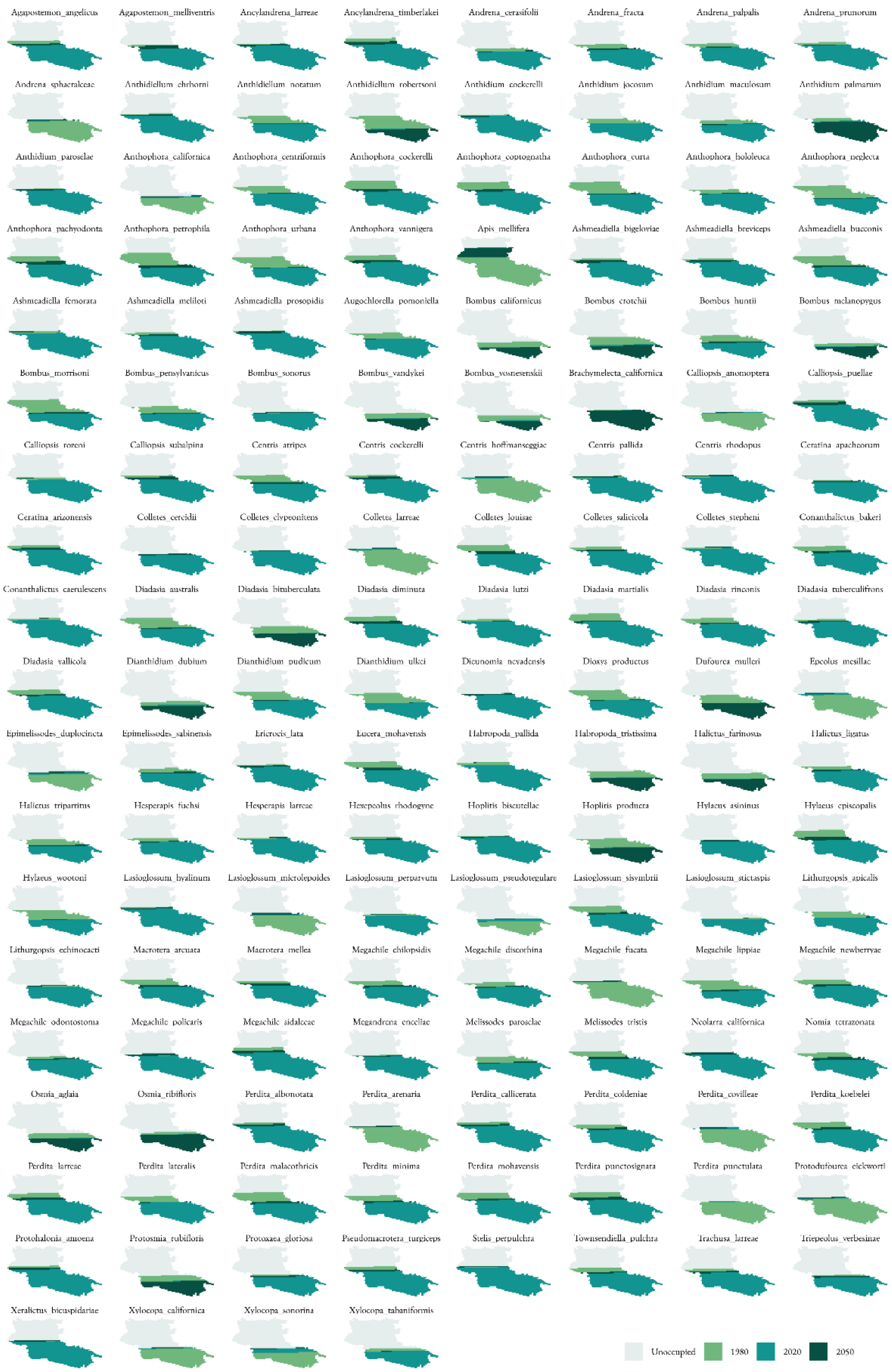


**Figure S1:** A representation of the range area of each of the modeled bee species. The colors indicate the proportion of the study area in which each species may occur during 1980, 2020, and 2050 under SSP 2-4.5.

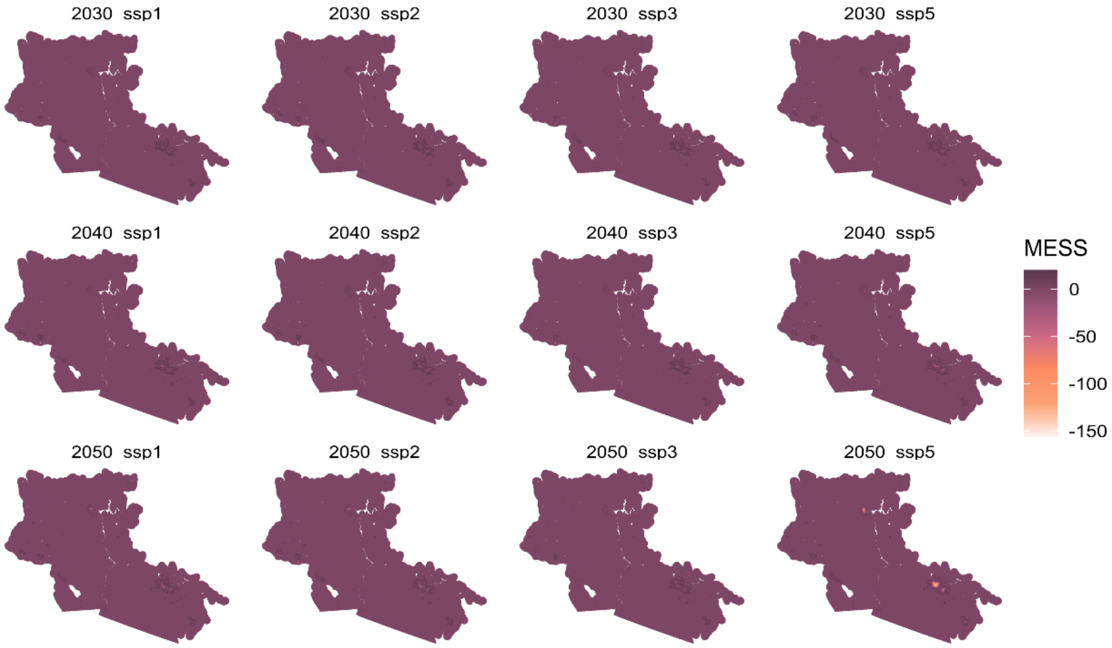


**Figure S2:** Multivariate Environmental Similarity Surfaces for each decade and climate change scenarios. Values less than 0 are novel environments compared to the training period. Novel environments are found in the urban cores of major metropolitan areas (>8 km from natural areas).

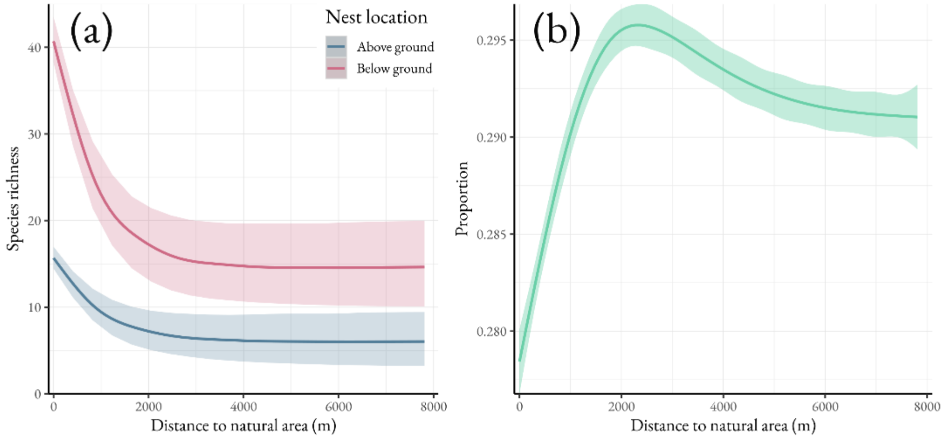


**Figure S3:** The species richness of above and below ground nesting bees across a gradient of distances from natural land uses within urban areas (a). The proportion of above ground nesting bees increases in urban areas (b).
